# Elongator is a microtubule polymerase selective for poly-glutamylated tubulin

**DOI:** 10.1101/2023.05.10.540202

**Authors:** Vicente José Planelles-Herrero, Mariya Genova, Alice Bittleston, Kerrie E. McNally, Gianluca Degliesposti, Maria M. Magiera, Carsten Janke, Emmanuel Derivery

## Abstract

Elongator is a tRNA-modifying complex that regulates the fidelity of protein translation. Recently, a moonlighting function of Elongator has been identified in regulating polarization of the microtubule cytoskeleton during asymmetric cell division. Elongator induces symmetry breaking of the anaphase midzone by selectively stabilizing microtubules on one side of the spindle. This polarizes the segregation of signalling endosomes containing cell-fate determinants to only one daughter cell, thus contributing to cell fate determination. Here, we unravelled the molecular mechanism by which Elongator controls microtubule dynamics. Elongator binds simultaneously to the tip of microtubules and also to free GTP-tubulin heterodimers via their C-terminal tails. Elongator thereby locally increases tubulin concentration at microtubule ends, which stabilizes microtubules by increasing their growth speed and decreasing their catastrophe rate. We show that the Elp123 and Elp456 subcomplexes bind to microtubules and free tubulin heterodimers, respectively, and that these activities must be coupled for Elongator to stabilize microtubules. Surprisingly, we found that Elp456 has strong selectivity towards polyglutamylated tubulin dimers. Hence, microtubules assembled by Elongator become selectively enriched with polyglutamylated tubulin. Therefore, Elongator can rewrite the tubulin code of growing microtubules, placing it at the core of cytoskeletal dynamics and polarization during asymmetric cell division.

## Introduction

Microtubules are dynamic cytoskeletal polymers found in every eukaryotic cell, where they are essential for cell division, morphogenesis, cell motility and intracellular transport. The structure, properties and dynamics of microtubules structures are tightly regulated by a plethora of proteins, including microtubule-associated proteins (MAPs), motor proteins and tubulin-modifying enzymes. Together, all these factors control the geometry of the microtubule landscape, leading to the formation of structures with highly distinctive shapes and behaviours, such as long and stable axonal microtubules, the dynamic mitotic spindle, or axonemal microtubules that mediate ciliary beating.

Microtubules are asymmetric hollow tubes built from heterodimers of *α*- and *β*-tubulin that are incorporated at both ends of the polymer: the slowly growing minus-end, and the fast growing plus-end. In cells, microtubule dynamics are primarily controlled by regulating the plus-end dynamics, while the minus-end is often anchored or protected from depolymerisation^1-3^. A particularly important group of proteins controlling microtubule dynamics is the so-called microtubule polymerases, which specifically recognize the growing-end of microtubules and increase their growth rate and/or decrease their catastrophe frequency^4-7^. Arguably the best characterized member of this family is XMAP215, a TOG (Tumour Overexpressed Gene)-domain containing protein. Through a series of tandem-linked TOG domains, XMAP215 polymerizes microtubules by simultaneously binding to the microtubule end and to free *αβ*-tubulin heterodimers, thus facilitating the integration of *αβ*-tubulin heterodimers into the growing microtubule^6,8^ On the other hand, CLASP, which contains only a single TOG domain, seems to stabilize microtubules using a different mechanism^7,9,10^. Indeed, a single, isolated TOG-domain has been shown to be sufficient to regulate microtubule polymerization dynamics. Finally, motor proteins have also been shown to modulate microtubule dynamics^5,7^. For example, kinesin-5, a tetrameric member of the kinesin family, enhances microtubule polymerization by stabilizing tubulin-tubulin interactions at the growing ends of microtubules^5^; whilst other kinesins, such as Kinesin-13, can actively depolymerise microtubules^11^.

Microtubules can also be modified by direct post-translational modifications of tubulin, further modulating their properties^12,13^. The C-terminal tails of both *α*- and *β*-tubulin are hotspots of modifications, in particular by tubulin tyrosine ligase-like (TTLL) proteins that add branching poly-glutamate peptide chains at several sites. This modification, known as poly-glutamylation, promotes the binding of several MAPs, or the action of microtubule-severing enzymes such as spastin and katanin. Poly-glutamylation has thus the potential to modulate microtubule dynamics and ultrastructure^14-19^. Since most, albeit not all, enzymes catalysing these post-translational modifications (PTMs) preferentially modify microtubules rather than soluble tubulin dimers^20-23^, it is currently thought that PTMs mostly change the behaviour of pre-existing microtubules.

The Elongator complex is a conserved molecular machine regulating protein translation. In particular, Elongator selectively modulates protein translation rates by modifying the wobble uridines at position 34 (U_34_) of a subset of tRNAs^24,25^. Structurally, Elongator is composed of six subunits, Elp1-6, which are each present in two copies and arranged in two discrete, stable sub-compiexes^26-29^: an Elp123 dimer and an Elp456 hexamer (**Fig. 1A**). In the context of its tRNA modification function, tRNAs bind to the larger, catalytically active Elp123 subcomplex ^24,25,28^. The function of the Elp456 sub-complex is less understood, but it is thought that its binding to the Elp123-tRNA complex releases the modified tRNA through a competition mechanism^28^.

**Figure 1.**
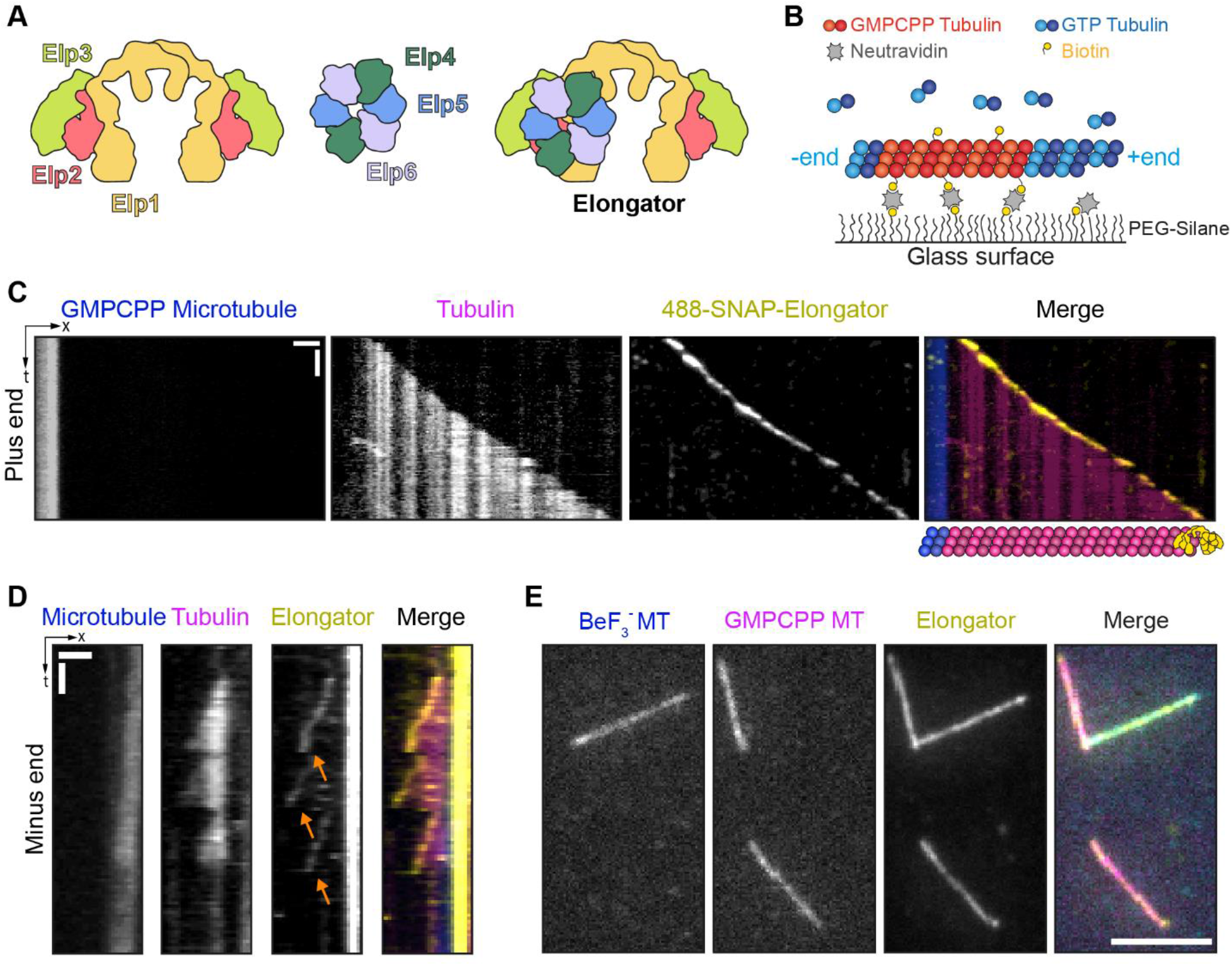
Elongator complex tracks the growing ends of microtubules. (**A**) Elongator contains one copy of two distinct stable sub-complexes: Elp123 (left) and the Elp456 ring (right). Note that although Elp123 contains two potential binding sites for Elp456, multiple assays confirmed that only one copy is preferentially bound^26,28,29^, making Elongator an asymmetric complex. (**B**) Assay design: Biotinylated, rhodamine-labelled GMPCPP-stabilized seeds (red) are anchored via NeutrAvidin to PLL-PEG-Silane. Free tubulin (17 μM 10% HiLyte 647-labelled, cyan) and 25 nM Alexa 488-SNAP-Elongator is added and microtubule polymerization is observed by TIRFM. (**C, D**) Representative kymographs showing Alexa488 SNAP-Elongator complex tracking the plus (C) and minus (D) ends of growing microtubules. Note that Elongator detaches from the microtubule end when microtubules undergo catastrophe (orange arrows). Also note Elongator accumulating at the transition point between the GMPCPP seed and the GDP lattice (see also. Sup. Fig. 1D). (**E**) Elongator decorates both GDP·BeF_3_ and GMPCPP stabilized microtubules. Scale bars = 2 min/2 μm (**C, D**), 5 μm (**E**).

Surprisingly, Elongator has recently been shown to control symmetry breaking of the anaphase spindle midzone during the asymmetric cell division of *Drosophila* Sensory Organ Precursor (SOP) cells^30^. This asymmetric central spindle in turns polarizes the segregation of signalling endosomes containing cell-fate determinants towards only one daughter cell, therefore contributing to asymmetric cell fate determination. Unexpectedly, Elongator’s activity of modulating microtubule dynamics was found to be independent on its effect on protein translation^30^. Rather, Elongator directly binds to microtubules and modulate their dynamics, specifically by increasing their growth rate and their lifetime by decreasing their catastrophe frequency^30^. Since Elongator is asymmetrically localized on one side of the anaphase spindle midzone, this induces preferential microtubule stabilization on one side of this spindle and thus symmetry breaking. However, how Elongator modulates microtubule dynamics at the molecular level was unknown.

Here, we reveal the molecular mechanism by which Elongator stabilizes microtubules. We show that Elongator can specifically recognise and track the growing ends of microtubules. By simultaneously binding to microtubule tips, via Elp123, and to *αβ*-tubulin heterodimers, via Elp456, Elongator increases the local concentration of tubulin dimers at growing ends, thereby increasing the growth rate and decreasing the catastrophe rate of microtubules. Strikingly, we show that Elp456 preferentially binds polyglutamylated tubulin heterodimers, while Elp123 binds to microtubules regardless of their post-translational modification. Thus, in the presence of Elongator, microtubules not only grow faster, but also get selectively and specifically enriched in poly-glutamylated tubulin. These results highlight an unexpected function for Elongator in remodelling the landscape of microtubule modifications, whilst itself not being a tubulin-modifying enzyme. Or work thus uncovers a novel molecular mechanism of how microtubule PTM diversity can be achieved in cells.

## Results

### Elongator is a microtubule end-tracking protein complex

To investigate the molecular mechanism by which Elongator binds and stabilizes microtubules, we capitalized on our previously described procedure for purifying the *Drosophila* complex from cultured S2 cells^30^ to derive an Alexa488-labelled SNAP-Elongator (**Fig. 1, Sup. Fig. S1A**). We then investigated the localization of this complex on dynamic microtubules using an established assay in which cycles of growth/catastrophe of dynamic microtubules from stable seeds are imaged by Total Internal Reflection Microscopy (TIRFM)^31^ (**Fig. 1B-D, Sup. Fig. S1B-C**). Under conditions in which Elongator and tubulin were at physiological concentrations (17 μM tubulin, 25 nM Alexa 488-SNAP-Elongator, ref.^32^) strong binding of Elongator to both the plus- (**Fig. 1C**) and minus-ends (**Fig. 1D**) of growing microtubules could be detected in ∼10% of the analysed growth events. The end-localization of Elongator was lost when the microtubule underwent catastrophe (**Fig. 1D, Sup. Fig. S1B-C**, orange arrows), showing that Elongator specifically recognizes the growing ends of microtubules. Moreover, and consistent with other end-tracking proteins^6,32,33^, Elongator could also be observed diffusing on the GMPCPP seeds (**Sup. Fig. S1B, C**), in conditions where Elongator tracks the growing end of microtubules (that is, when free tubulin is present). Strikingly, in these conditions, Elongator could be observed moving from the seed to the growing end of a microtubule (**Sup. Fig. S1C**, yellow arrows), highlighting its preference for GTP-like structures. Consistently, the GDP-lattice of the microtubule was mostly devoid of Elongator signal (**Fig. 1C,D**).

The growing end of microtubules (also known as the microtubule “tip” or “cap”) is formed by newly incorporated GTP-*αβ*-tubulin heterodimers. Once incorporated, GTP is hydrolysed to GDP, creating a characteristic GDP microtubule shaft with a “GTP-cap” located at the growing end of the polymer. To better understand how Elongator binds to the end of microtubules, we examined the binding of Elongator to GTP-mimicking microtubules. First, we used the GTP analogue GMPCPP to grow microtubules, which mimics key features of the GTP-state of microtubule ends^34,35^. As we previously reported^30^, Elongator binds and decorates GMPCPP-stabilized microtubules along their entire length (**Fig. 1E**). To confirm that tip-recognition involved recognition of the tubulin nucleotide state, we used BeF_3_^-^ microtubule seeds, which are known to more precisely mimic the GTP-state of the microtubules than GMPCPP^34^. Elongator was consistently found to bind BeF_3_^-^-bound stabilized microtubules with similar apparent affinities (**Fig. 1E**). This establishes that Elongator specifically recognizes the GTP (or GTP-like) state of the growing ends of microtubules, both at the minus and at the plus ends. Note that an increased Elongator signal can also be observed at transition points between the GMPCPP seed and the GDP lattice (**Fig. 1D, Sup Fig. S1D**), a feature seen in end-binding (EB) proteins^33^, suggesting that the binding to GTP-cap could be facilitated by the open, tapered microtubule ends.

### The Elp123 subcomplex binds to, but does not stabilise, microtubules

We next investigated how Elongator binds to microtubules. An isolated Elp2 subunit produced in yeast has been shown to interact with microtubules at high concentration^36^, and we previously reported that neither Elp456 hexamers nor a partial subcomplex of Elp23 with interacting protein Hsc70-4 exhibit appreciable microtubule binding^30^. Since the full Elongator complex, consisting of two copies of Elp123, dimerizing *via* Elp1, and one Elp456 hexameric ring (**Fig. 1A**) binds to microtubules at very low concentrations (**Fig. 1**), we speculated that the full, dimeric Elp123 sub-complex might be needed for this activity.

To purify the Elp123 subcomplex, we used the biG-Bac expression system to simultaneously overexpress *Drosophila* Elp1,2 and 3 in Sf9 insect cells^37^(**Sup. Fig. S2A-C**). We tagged Elp3 since it is the only subunit in fly Elp123 that can accommodate a SNAP-tag whilst being compatible with the assembly of the full complex^30^. After two affinity purification steps, the gel filtration profile shows that a highly pure Elp123 elutes at the expected elution volume (∼600 kDa, **Sup. Fig. S2B**), showing that *Drosophila* Elp123 dimerizes as expected^26,28,29^.

Strikingly we found that Alexa 488-SNAP-Elp123 directly bind to microtubules using TIRFM (**Fig. 2A, Sup. Fig. S2F** left panel) and microtubule pelleting assays (**Sup. Fig. S2D**), confirming that the Elp123 subcomplex is sufficient for microtubule binding. Yet, Elp123 does not stabilize microtubules (**Fig. 2B, C**). Indeed, using the same microtubule dynamics assays depicted before (**Fig. 1B**), no effect was detected in the microtubule growth speed (**Fig. 2B**, mean ± SEM: 1.003 ± 0.009 μm/min for the buffer control, versus 1.050 ± 0.009 μm/min for Elp123, n=458) or lifetime (**Fig. 2C**, 202.61 ± 7.10 s versus 211.76 ± 8.14 s) at the plus end. This is in sharp contrast to the effects of the full Elongator complex at the plus end in similar conditions, which shows a 1.4-fold increase of the growth speed (dashed line in **Fig. 2C**) and a 1.8-fold increase of microtubule lifetime^30^. Similarly, we also could not detect any effect of Elp123 at the minus ends (control: 0.319 ± 0.007 μm/min, n=90: Elp123:0.343 ± 0.010 μm/min, n=138)^24^.

**Figure 2.**
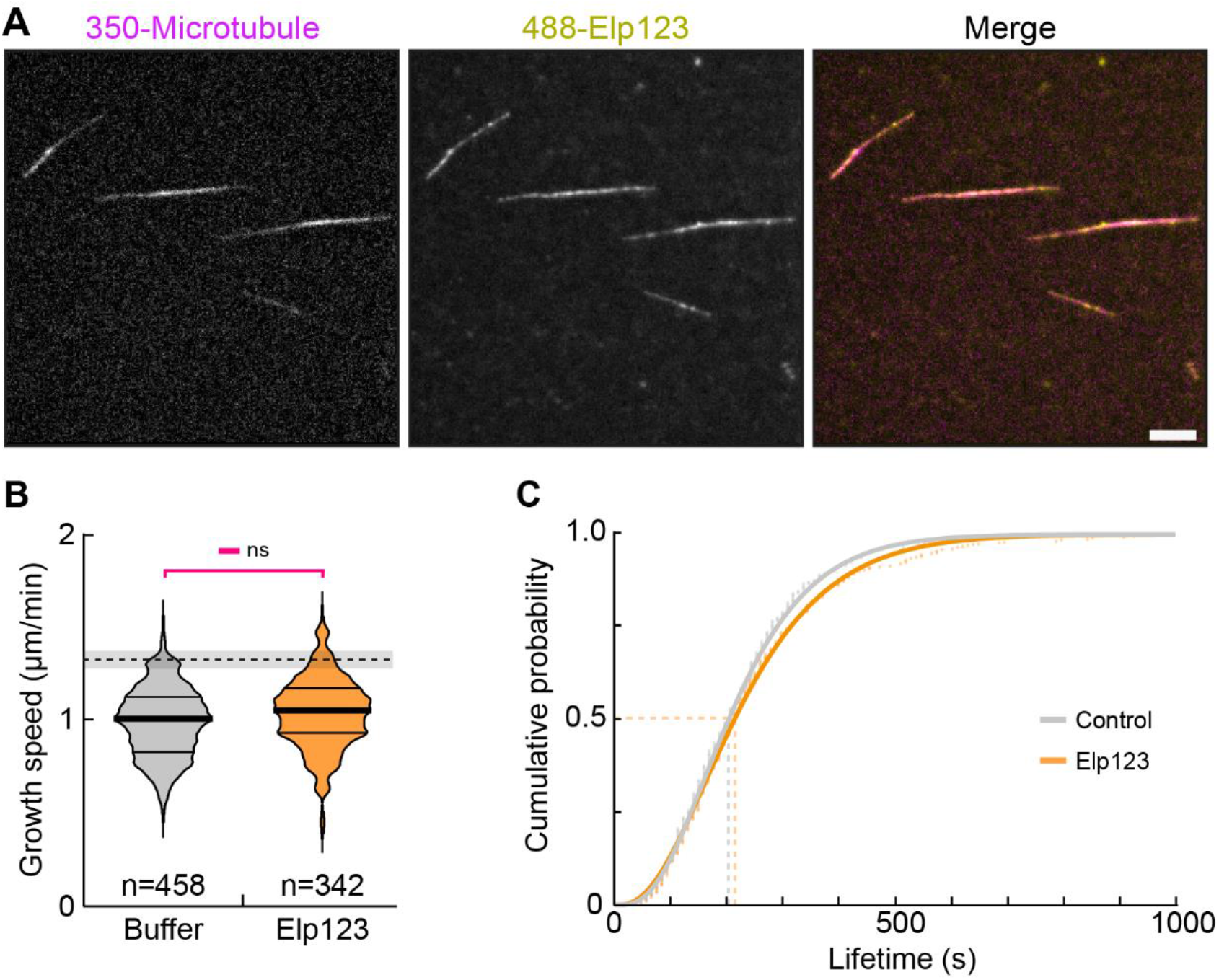
Elp123 binds but does not stabilize microtubules. (**A**) AMCA-labelled GMPCPP stabilized microtubules incubated with 25 nM Elp123 sub-complex labelled with AlexaFluor488 dye and observed by TIRFM. Elp123 directly binds to GMPCPP microtubules. (**B**) Microtubule seeds were polymerized in the presence of 16 μM GTP-tubulin (10% HiLyte 647 labelled) in the absence or presence of 25 nM (His)_6_-PC-SNAP-Elp123, and microtubule growth rate at the plus end was measured by TIRFM. The control was performed using the same buffer as Elongator kept from the final purification step (see Sup. Fig. S2A). P value from a two-sided Kruskal-Wallis test for non-parametric samples is indicated; n, number of individual growth events quantified from at least three independent experiments. Dashed line represents an increase of ∼1.4 fold in the speed of microtubule growth, which happens in the presence of the full Elongator complex^30^. Thick line, median; thin line, quartile. (**C**) Cumulative microtubule lifetime distribution of microtubules grown at the plus end in the absence and presence of Elp123. Elp123 does not stabilize microtubules. Mean lifetime estimate ± error (lifetime at half cumulative distribution, dashed lines): 202.61 ± 7.10 s (Control) and 211.76 ± 8.14 s (Elp123). Lines, gamma distribution fits. Scale bar = 5 μm.

Altogether, our data show that, although Elp123 directly interacts with microtubules (**Fig. 2A**), this is not sufficient to modulate tubulin dynamics, in particular microtubule stabilization. Thus, an as-of-yet uncharacterized mechanism must be responsible for this effect. Since the Elp456 sub-complex was absent in this experiment, we examined the role of Elp456 in microtubule stabilization.

### Elp456 binds to *αβ*-tubulin heterodimers

We noticed in our reconstitution of microtubule dynamics in the presence of the full Elongator complex that HiLyte 647-*αβ*-tubulin signal could be seen diffusing together with Alexa488-SNAP-Elongator on microtubules (**Sup. Fig. S1C**, magenta arrows). This suggested that Elongator might be able to interact simultaneously with both microtubules and soluble *αβ*-tubulin heterodimers, as previously shown for TOG-domain containing proteins^610^. To verify this, we added the different proteins into our reconstitution assay sequentially, as opposed to simultaneously. We first incubated GMPCPP-stabilized microtubules with Alexa 488-SNAP-Elongator. Then, an excess of Hilyte 647-*αβ*-tubuin was added and the chamber was imaged by TIRFM (**Fig. 3A**). We found that Hilyte 647-*αβ*-tubulin signal could be observed decorating microtubules, but only in the presence of Elongator (**Fig. 3A**). This establishes that Elongator can interact with both microtubules and tubulin.

**Figure 3.**
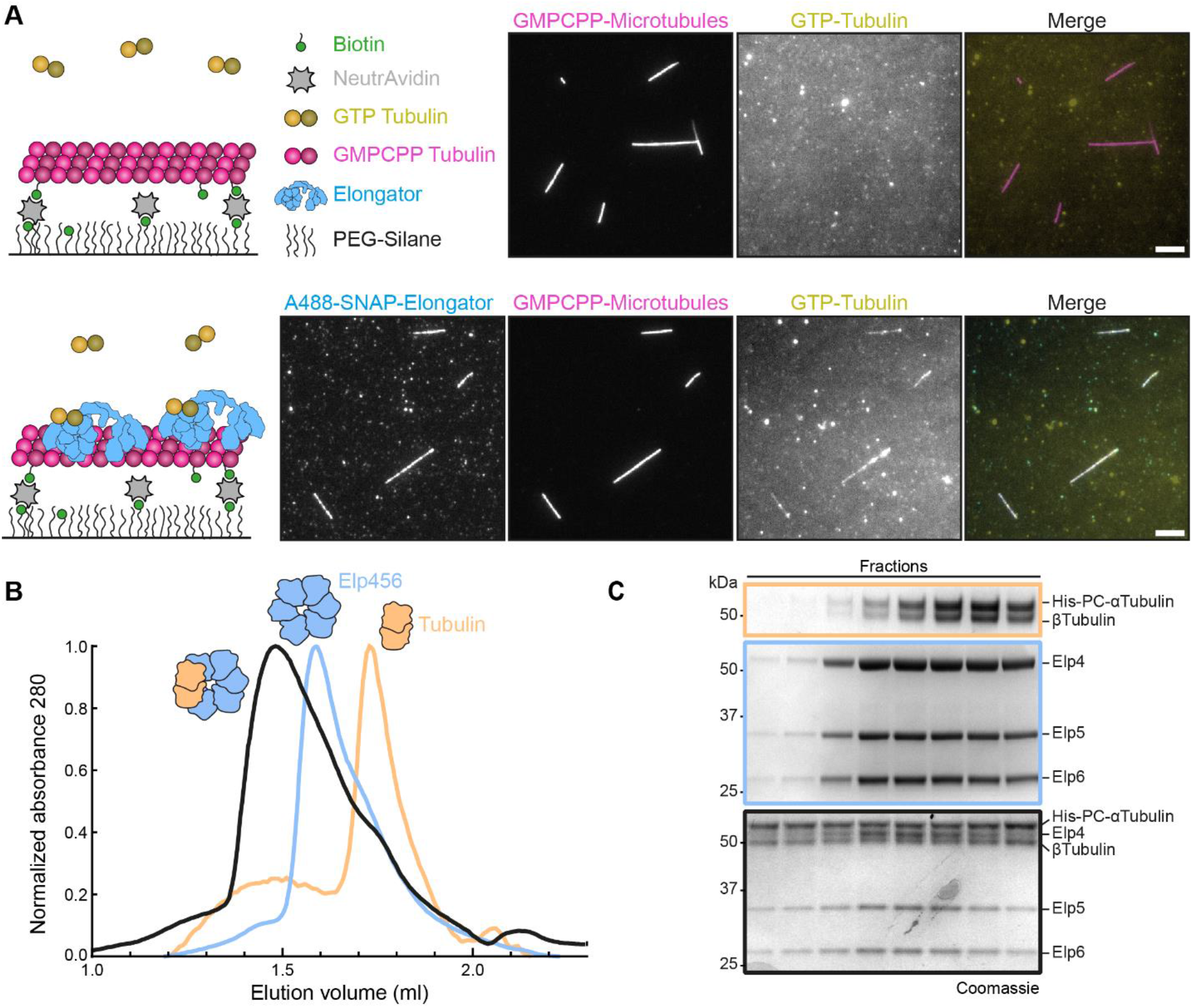
Elongator complex binds to tubulin through Elp456. (**A**) While bound to GMPCPP-stabilized microtubules, the Elongator complex can recruit additional globular GTP-tubulin. Top row: preformed rhodamine-labelled. GMPCPP-stabilized microtubules incubated with HiLytc647-labelled GTP-tubulin (16.7 µM GTP-tubulin, 10% HiLytc647-tubulin). Bottom row: preformed Rhodaminc-labelled, GMPCPP-stabilized microtubules incubated first with AlexaFluor488-SNAP-Elongator complex, then with HiLyte647-labelled GTP-tubulin as above. (**B**) Elp456 binds to recombinant *Drosophila α*1*β*1-tubulin heterodimers. Size-exclusion elution profiles of tubulin (orange), Elp456 (blue) and a mixture containing both proteins in a 1.2:l ratio (tubulin:Elp456, black). The Elp456-tubulin complex elutes earlier from a superpose 6 increase 3.2/300 column. (**C**) All elution profiles shown in (**B**) were analysed by Coomassie blue, confirming a shift in the protein elution patterns. Due to the presence of a (His)_6_-PC tag in *α*tubulin a difference in size can be observed between (His)_6_-PC-α and *β*-tubulin. Scale bars = 5 µm.

However, Elongator does not contain any TOG-domain, or other sequence motives predicted to interact with unpolymerized, *αβ*-tubulin heterodimers. We thus wondered if Elp456, whose role in Elongator function is still not fully understood^28,38^, could bind to tubulin using as-of-yet uncharacterized surfaces. For this, we purified the *Drosophila* Elp456 subcomplex from *E. coli* (**Sup. Fig. S3A, B**) and investigated its interactions with tubulin. Importantly, using size-exclusion chromatography, we found that *Drosophila* Elp456 binds to recombinant *Drosophila* α1β1-tubulin heterodimers (**Fig. 3B, C**) in solution in a seemingly 1:1 ratio (Elp456 hexameric ring:tubulin dimer). We confirmed this result by immunoprecipitation using the affinity tag present on our recombinant tubulin (**Sup. Fig. S3C**). To determine if the binding of *αβ*-tubulin heterodimers was specific to Elp456, we assessed Elp123 ability to bind tubulin under similar conditions. Critically, although Elp123 did bind to microtubules (**Fig. 2A, Sup. Fig. S2D**), it did not bind to *αβ*-tubulin heterodimers (**Sup. Fig. S3D**).

All together, these results reveal a clear separation of function for the two Elongator sub-complexes: Elp123 binds to microtubules (**Fig. 2A**), but not to free (GTP-) tubulin (**Sup. Fig. S3D**), and, conversely, Elp456 binds to free tubulin (**Fig. 3B**) but not microtubules^30^.

### Elongator binding to both microtubules and tubulin is needed for microtubule stabilization

We reasoned that the effect of Elongator on microtubule stabilization must require both the microtubule- and the *αβ*-tubulin-binding moieties, since, unlike the full Elongator complex, neither Elp123 (**Fig. 2**) nor Elp456 (**Fig. 4B**) on their own showed any effect on microtubule dynamics. We sought to test this hypothesis by investigating the effects on microtubule dynamics of a reconstituted full Elongator, which was generated by mixing the Elp123 and Elp456 subcomplexes together after purification (**Fig. 4**).

**Figure 4.**
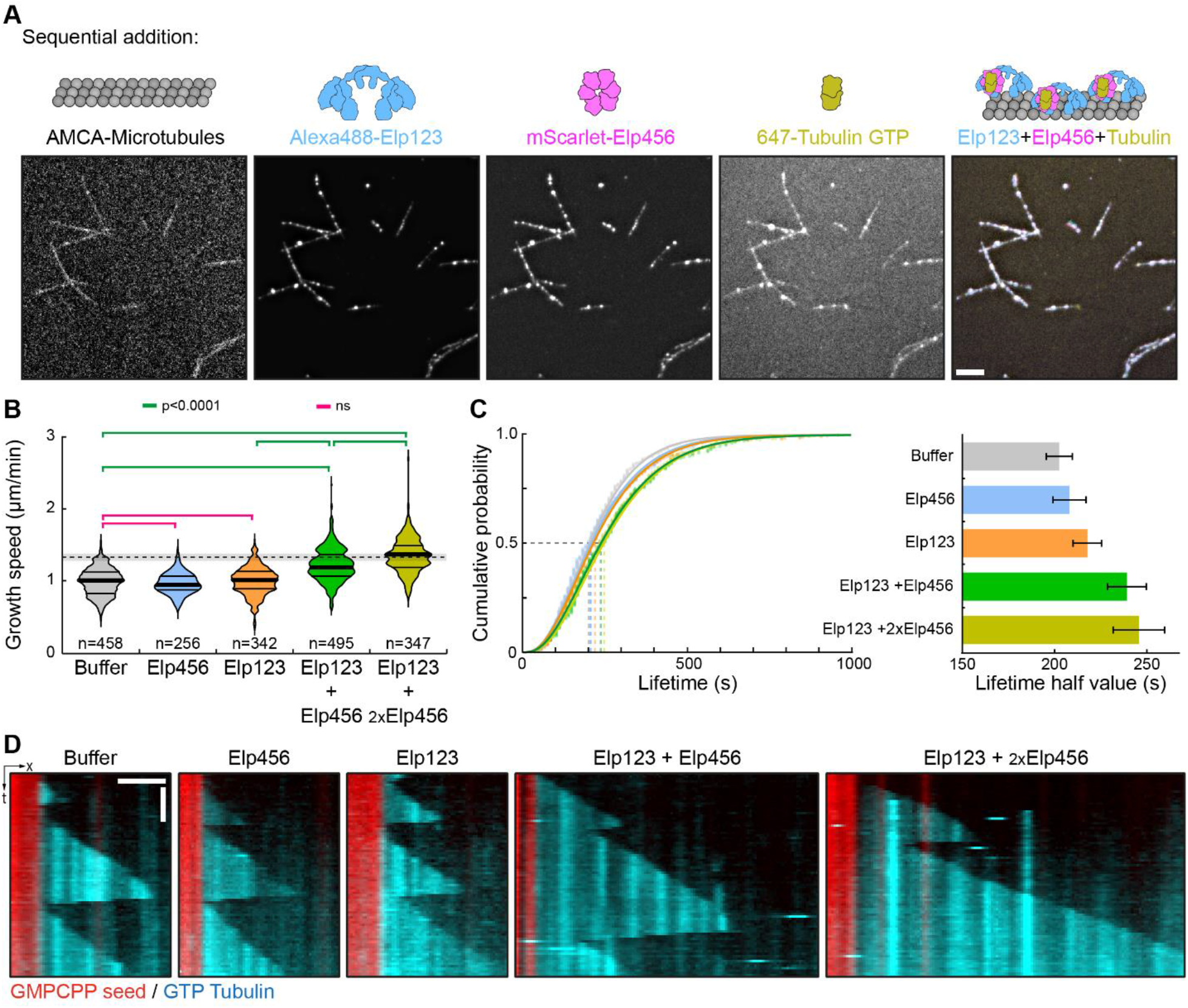
Binding to both microtubules and free tubulin is essential for microtubule stabilization. (**A**) Sequential reconstitution of a full Elongator complex onto microtubules using independently-purified Elp123 and Elp456 subcomplexes. Indicated components were sequentially added starting with AMCA-labelled microtubules, followed by AlexaFluor488-SNAP-Elp123, mScarlet-Elp456 and finally HiLyte647-tubulin. Note that all three proteins are observed on microtubules, then AlexaFluor488-SNAP-Elp123, then mScarlet-Elp456 and finally HiLyte647-tubulin. Note that all three proteins can be observed on microtubules. See Sup. Fig. S42A for drop out controls. For visualisation purposes, images were processed with a Wavelet “à trous” denoising filter. (**B, C**) Effect of the indicated conditions on the growth speed (B) and lifetime (C) of microtubules at the plus end imaged by TIRFM in the presence of 16 µM GTP-tubulin (10% HiLyte 647 labelled). Elongator concentrations are: 100 nM Elp456 (“Elp456”), 50 nM (His)_6_-PC-SNAP-Elp123 (“Elp123”), 50 nM Elp456+50 nM (His)_6_-PC-SNAP-Elp123 (“Elp123+Elp456”), and 100 nM Elp456+50 nM (His)_6_-PC-SNAP-Elp123 (“Elp123+2xElp456”). n, number of micro tubule-growing events analysed. (**B**) P values for a Kruskal-Wallis test followed by Dunn’s multiple comparison test are indicated. Dashed line represents an increase of ∼1.4 in the speed of microtubule growth^30^. Thick line, median; thin line, quartile. (**C**) Microtubule lifetime estimate ± error from the bootstrapped mean lifetimes (see methods) are indicated in the right panel. (**D**) Representative kymographs of the indicated conditions. Scale bars = 5 μm (**A**), 2 min/2μm (**D**).

We first confirmed that the full Elongator complex can be reconstituted on GMPCPP microtubules by sequentially adding Elp123 then Elp456 subcomplexes, and, more importantly, that the reconstituted complex is functional as assessed by its ability to capture *αβ*-tubulin heterodimer once bound to the microtubule lattice (**Fig. 4A**). For this, we sequentially added in the same chamber fluorescently labelled AMCA-microtubules, 488-SNAP-Elp123, mScarlet-Elp456 followed by Alexa647-*αβ*-tubulin. Strikingly, tubulin heterodimers could be recruited onto microtubules by the reconstituted full Elongator (i.e. Elp123+Elp456, see **Fig. 4A** and **Sup. Fig. S4** for drop out controls), which therefore mimics the behaviour observed with the native full Elongator complex purified from *Drosophila* cells (**Fig. 3A**). As expected, in the absence of Elp456, tubulin was not recruited to microtubules (**Sup. Fig. S4A**), in line with the fact that Elp123 alone does not bind to tubulin (**Sup. Fig. S3D**). Note that at the high concentrations used in this experiment, a very weak mScarlet-Elp456 signal could be detected on microtubules, but this was markedly less seen than in the presence of Elp123 (9.3-fold less fluorescence intensity, **Sup. Fig. S4A**, third versus first row, **Sup. Fig. S4B** for quantification). Together, these results confirm that the full Elongator can bind simultaneously to both microtubules and *αβ*-tubulin heterodimers via Elp123 and Elp456, respectively.

We next investigated the effects of the reconstituted Elongator complex on microtubules dynamics. Remarkably, when both Elp123 and Elp456 were added to the assay, microtubules grew significantly faster compared to the buffer alone or the two individual sub-complexes (**Fig. 4B, D**). In the presence of reconstituted Elongator, microtubule growth speed increases ∼1.35-fold at the plus end in the presence of Elp123 and an excess of Elp456 (1.020 ± 0.009 μm/min to 1.380 ± 0.013 μm/min), which is remarkably similar to the ∼1.4-fold increase we reported for the full Elongator complex purified from *Drosophila* cells^30^. A similar effect was observed at the minus-ends, again matching the ∼1.2 fold increase we reported for the full Elongator complex^30^ (control: 0.319 ± 0.007 μm/min, n=90; Elp456:0.329 ±0.011 μm/min, n=72; Elp123:0.343 ± 0.010 μm/min, n=138; Elp123+Elp456: 0.389 ± 0.010 μm/min, n=133; Elp123+2xElp456:0.420 ± 0.014 μm/min, n=100). Furthermore, the microtubule lifetime was increased in the presence of both sub-complexes (**Fig. 4C**), again recapitulating the effect observed with the full Elongator complex^30^. Altogether, these results suggest that the effect of Elongator on microtubule dynamics require both its microtubule and tubulin heterodimer binding activities via its Elp123 and Elp456 subcomplexes, respectively.

### Coordinated action between subcomplexes is critical for microtubule stabilization

We showed that Elp123 binds to microtubules (**Fig. 2, Sup. Fig. S2**), Elp456 binds to *αβ*-tubulin heterodimers (**Fig. 3B, Sup. Fig. S3C**) and that both activities are required to reconstitute the activity of the full complex (**Fig. 4**). Since Elongator is recruited to the growing ends of microtubules (**Fig. 1**), this suggests a mechanism whereby Elongator would increase microtubule growth speed by bringing additional free *αβ*-tubulin heterodimers close to the tip of the microtubules. This would increase the local concentration of tubulin at the growing ends, thereby increasing the growth speed of microtubules and decreasing their catastrophe rate, which could explain Elongator’s effects on microtubule dynamics (**Fig. 5A**, left). Elongator would thus stabilize microtubules in a mechanism reminiscent of microtubule polymerases such as XMAP215 and CLASPs^6,8,10^. Importantly, we showed that the Elp456-*αβ*-tubulin pre-formed complex could be recruited to Elp123 already bound to microtubules (**Sup. Fig. S4C**), further supporting this mechanism of action. We thus thought to probe the importance of the coupling between the activities of Elp123 and Elp456 for the effect of the full complex on microtubules (**Fig. 5**).

**Figure 5.**
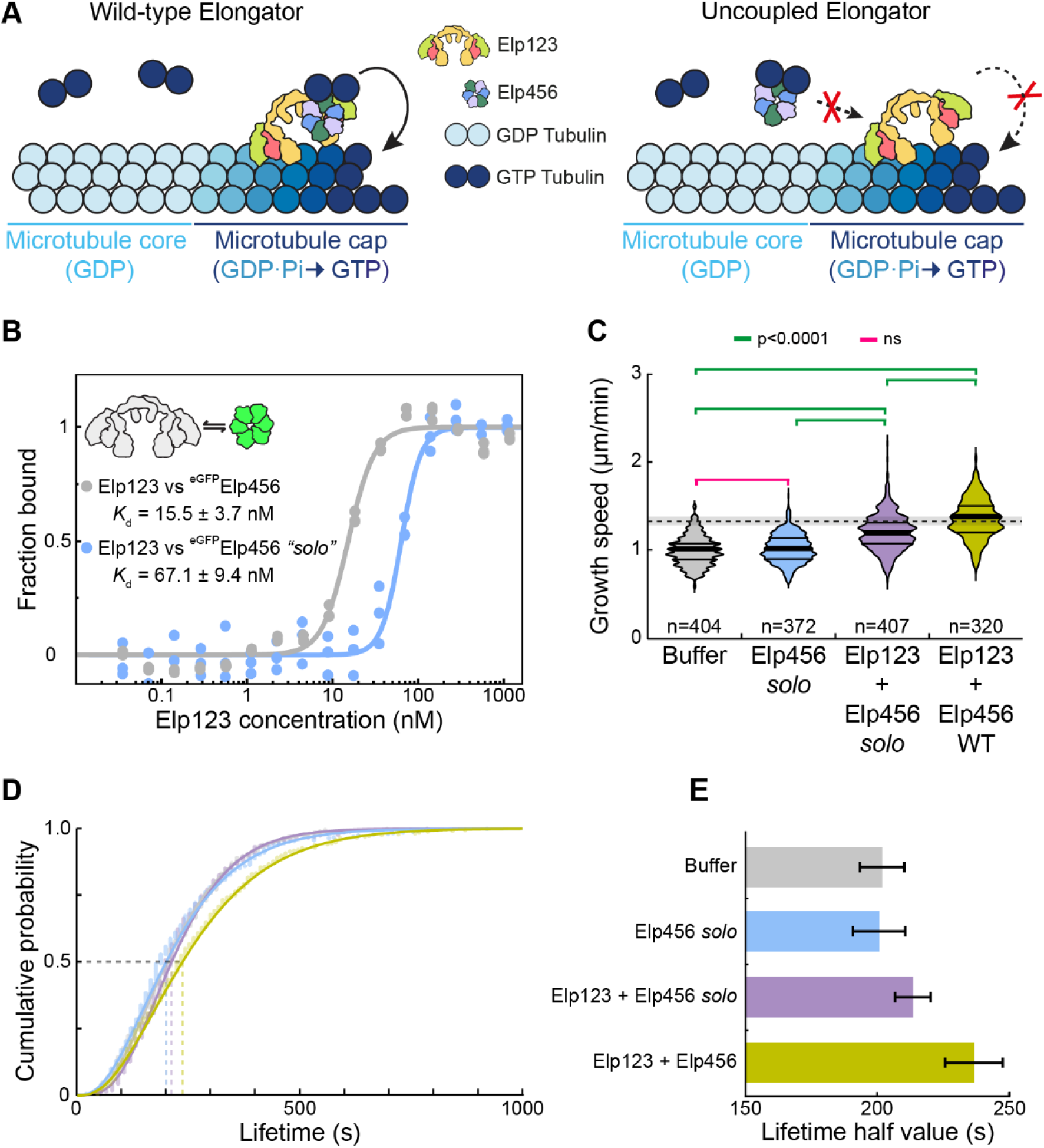
Elp123 and Elp456 coordinated action is critical for microtubule stabilization. (**A**) Proposed molecular mechanism of Elongator’s mode of action on microtubules. Elongator binds to the microtubule tips via Elp123 and to *α β*-tubulin heterodimers via Elp456. This increases the local tubulin concentration at the microtubule tips, thereby increasing the polymerization speed and decreasing the catastrophe rate. If the interaction between Elp123 and Elp456 is perturbed, the effect on microtubule dynamics is reduced. (**B**) Measurement of the Elp123 and eGFP-Elp456 WT and *solo* mutant interaction using microscale thermophoresis (see methods). Calculated dissociation constant (*K*_d_) values are indicated (mean ± s.d.; n=3). The calculated *K*_d_ for the Elp456 *solo* mutant is 4.3 times lower than for the wild-type. (**C, D, E**) Effect of the indicated proteins on the growth speed (C) and lifetime (D, E) of microtubules at the plus end measured by TIRFM in the presence of 16 µM GTP-tubulin (10% HiLyte 647 labelled). Elongator concentrations are: 100 nM Elp456 *solo* (“Elp456 solo”), 100 nM Elp456 *solo* +50 nM (His)_6_-PC-SNAP-Elp123 (“Elp123+Elp456 solo”), and 100 nM Elp456 wild-type+50 nM (His)_6_-PC-SNAP-Elp123 (“Elp123+Elp456”). n, number of microtubule-growing events analysed. (**C**) P values for a Kruskal-Wallis test followed by Dunn’s multiple comparison test are indicated. Dashed line represents an increase of ∼l.4 in the speed of microtubule growth^30^. Thick line, median; thin line, quartile. (**E**) Microtubule lifetime estimate ± error from the bootstrapped mean lifetimes (see methods).

We sought to produce a mutant of Elp456 displaying weaker affinity towards Elp123, whilst maintaining full *αβ*-tubulin binding capacity, resulting in uncoupling between the activities of the two Elongator sub-complexes (**Fig. 5A**, right). We first performed cross-linking mass spectrometry on the full Elongator complex to identify residues at the interface between *Drosophila* Elp123 and Elp456(see methods). This approach yielded several fragments containing cross-linked lysine residues (**Sup. Fig. S5A-C, E, Sup. Table S1**). We then cross-referenced these results with the localization of various Elongator disease-related mutations^39^ (**Sup. Fig. S5E**), conservation analysis (**Sup. Fig. S5A-C**), and AlphaFold2 Multimer^40,41^ modelling (see methods) to infer the surfaces of Elp456 involved in Elp123 binding (**Sup. Fig. S5E**). This allowed the rational design of a mutant harbouring 3-point mutations in each subunit (18 point mutations per Elp456 hexameric ring: Elp4 K364E, R397E, E410R; Elp5 K151E, K179E, T181A; Elp6 K119E, S200W, K228E; **Sup. Fig. S5A-C, F**). Importantly, we made sure to preserve the solvent-exposed surface of Elp456 (i.e., opposite to the surface that binds to Elp123) to assure that this this mutant could retain full binding to *αβ*-tubulin heterodimers. Hereafter, we refer to this mutant as Elp456 “solo” due to its predicted weaker binding to Elp123.

Elp456 *solo* could be readily purified from bacteria using the same protocol as for *wild-type* Elp456 (see methods). We then characterized the biochemical properties of Elp456 *solo* (**Fig. 5, Sup. Fig. S5**), namely its ability to bind Elp123 and to *αβ*-tubuIin heterodimers. First, we verified using cryo-electron microscopy (CryoEM) that the overall structure of Elp456 is not perturbed by the point mutations (**Sup. Fig. S5D**). Then we confirmed that the binding to recombinant *Drosophila* tubulin is not compromised in the *solo* mutant (*K*_d_; 1.6 ± 0.5 μM vs *wild-type* 1.2 ± 0.8 μM, **Sup. Fig. S5G**). Critically, however, the binding of Elp456 solo to Elp123 was significantly reduced compared to the *wild-type* (>4-fold reduction, from *K*_d_ of 15.5 ± 3.7 nM to 67.1 ± 9.4 nM, **Fig. 5B**). Importantly, the measured *K*_d_s are in the range of protein concentrations used in our *in vitro* experiments (i.e., 50 nM), thus we do expect a difference of occupancy in our assays. Indeed, at the concentrations used (i.e., 50 nM of Elp123 and 100 nM of Elp456), a ∼35% reduction in the total amount of Elp456 bound to Elp123 is expected (see methods). We then verified this prediction by using the Elp456 *solo* mutant in microtubule dynamics assays (**Fig. 5C-E**). Strikingly, the reconstituted Elongator containing Elp456 *solo* had an intermediate effect between the buffer alone and reconstituted Elongator containing *wild type* Elp456, both for the microtubule growth speed (**Fig. 5C**) and the lifetime (**Fig. 5D, E**) of microtubules. These intermediate, rather than total, effects are consistent with the relative affinity of Elp456 *solo* towards Elp123 (**Fig. 5B**). Furthermore, the somewhat mild affinity of wild-type Elp456 for Elp123 (15.5 nM) also explains why, in our conditions, we could detect a dose-dependence effect when increasing amounts of *wild type* Elp456 were added to Elp123 (**Fig. 4B, C, D**). Note that a recently published structure of the full S. *cerevisiae* and *M. musculus* Elongator complexes^28^ confirmed that most of the residues mutated in *D. melanogaster* Elp456 *solo* are indeed expected to be involved in the binding to Elp123, validating our predictions. For example, the Elp6 K199E and S200W mutations lie at the interface between Elp3 and Elp6 (**Sup. Fig. S5F** inset), and the introduction of a negative charge and a bulky tryptophan, respectively, are expected to greatly perturb this interface.

Altogether, our results establish that a tight coupling between both sub-complexes is required for Elongator’s ability to stabilize microtubules, whereby the Elp456-*αβ*-tubulin complex binds to Elp123 and thereby increase the local concentration of tubulin in the vicinity of the microtubule tip (**Fig. 5A**).

### Elongator can discriminate tubulin with specific post-translational modifications

We next investigated how Elongator can recognise and recruit free *αβ*-tubulin dimers at the molecular level, since Elp456 did not contain any predicted tubulin- (or microtubule-) binding surfaces. We thus decided to analyse the stoichiometric complex between Elp456 and tubulin by CryoEM to gain insights into the surfaces involved in the binding. However, although a very stable complex between Elp456 and tubulin can be isolated (**Fig. 3B, C, Sup. Fig. S3B,C**), tubulin could never be observed as extra signal compared to the control (**Sup. Fig. S6A**), even in cross-linked samples and following various strategies to maximize the signal outside of the Elp456 ring (see methods). We thus hypothesized that the binding must involve very flexible regions of Elp456, tubulin, or both, which could explain the absence of extra density for tubulin after averaging (see methods).

We thus inspected more carefully Elp456 and found that *Drosophila melanogoster* Elp456 harbours a specific, proline- and asparagine-rich insert in Elp4 (**Sup. Fig. S7**), which is predicted to be flexible and exposed to the solvent (Sup. Fig S7B, C). This insert connectsan *α*-helix and a *β*-strand that are usually held together by a short linker, as seen in several yeast Elp456 structures (**Sup. Fig. S7A, C**), and contains long stretches of polar residues surrounded by either negative or positive charges (**Sup. Fig. S7A**). Since tubulin is negatively charged, this *Drosophila* Elp456 linker could have been involved in the binding, so we removed it and purified the resulting Elp456 ΔpolyN mutant (**Sup. Fig. S7D**). The reconstituted Elp123+Elp456-ΔpolyN completely recapitulates the effect measured with wild-type Elp456 both for the microtubule growth speed (**Sup. Fig. S7E**) and lifetime (**Sup. Fig. S7F, G**) in TIRFM assays, confirming that this flexible region is not required for tubulin binding. We also observed that Elp5 contains a very flexible and disordered region at its C-termini (**Sup. Fig. S7B**). However, this region is very negatively charged and its participation in the binding with tubulin is therefore unlikely

We thus focused on the interaction partner tubulin. All tubulins contain a very characteristic stretch of disordered residues at the C-terminus, known as C-terminal “tails”. These tails extend from the tubulin body and are known to regulate the intrinsic properties of microtubules, as well as regulating the binding to associated proteins notably using a plethora of post-translational modifications (PTMs)^42-47^, Subtilisin-digestion of polymerized microtubules is an established way to specifically remove the C-terminal tails of tubulin. However, since in our hands subtilisin-digested tubulin is not stable when depolymerized, presumably due to the multiple digestion sites present in depolymerized tubulin^48^, we thought to use tubulins lacking post-translational modifications due to the specific source they are purified from (see methods). We first tested the binding of Elp456 against tubulin purified from HeLa S3 cells, known to be almost entirely free of poly-glutamylation, acetylation and de- tyrosination (i.e. non-modified)^49,50^. Strikingly, we detected an ∼18-fold reduction in the affinity of Elp456 towards HeLa S3 tubulin (*K*_d_: 21.2 ± 7.8 μM, n=3) compared to tubulin purified from mouse (*K*_d_: 1.8 ± 0.6 μM, n=3), pig brains (*K*_d_: 1.8 ± 0.5 μM, n=3) or wild-type, iso type-pure. *Drosophila* recombinant α1β1-tubulin (*K*_d_: 1.2 ± 0.8 μM, n=3) (**Fig. 6A-B, Sup. Fig. S6B, Sup. Fig. S6C**).

**Figure 6.**
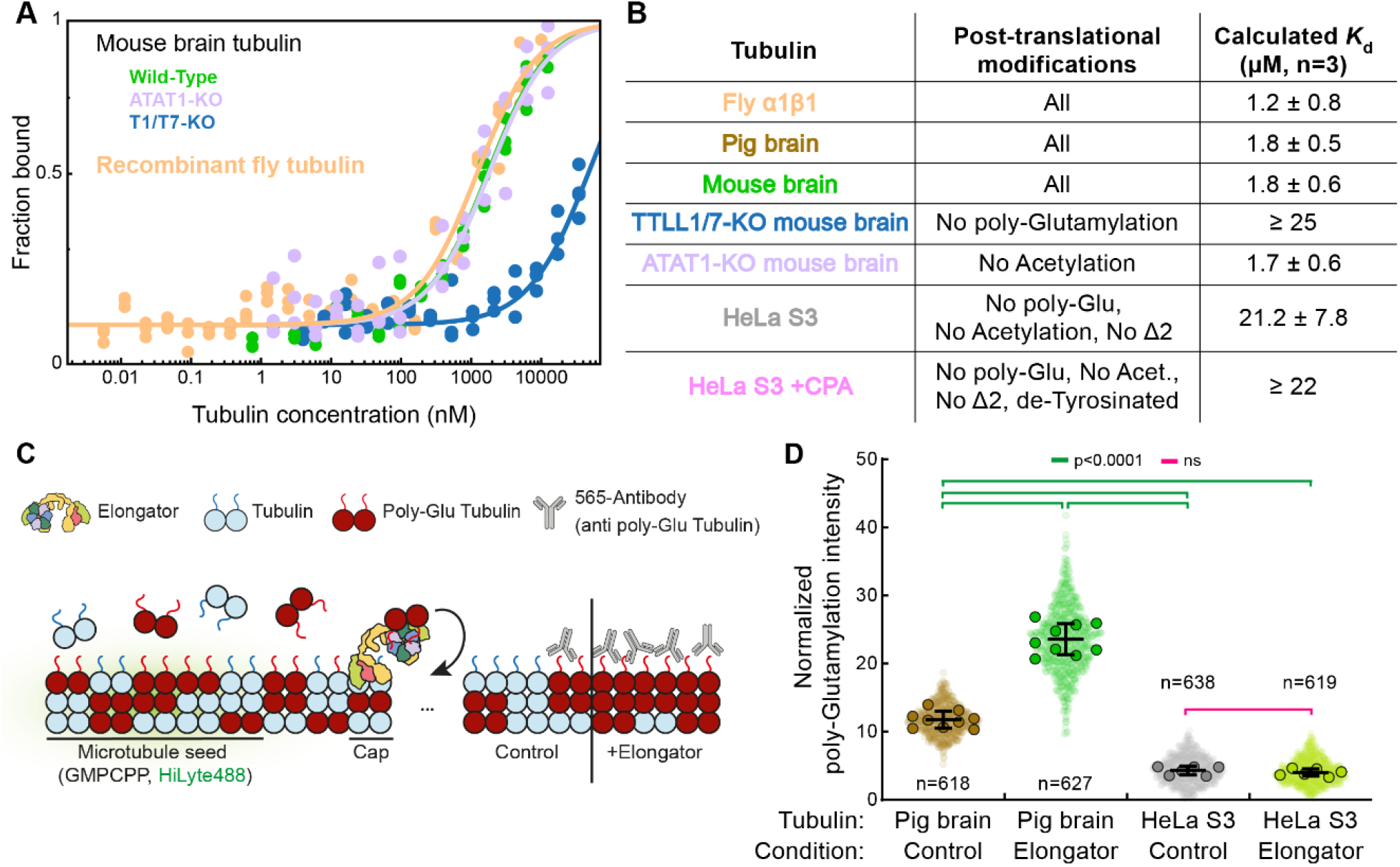
Elongator is a tubulin polymerase selective for poly-glutamylated monomers. (**A, B**) Elp456 binds preferentially to poly-glutamylated tubulin. (**A**) Affinity measurement between eGFP-Elp456 and indicated *αβ* -tubulin heterodimers using microscale thermophoresis (see methods). Calculated dissociation constant (*K*_d_) values are indicated (mean ± s.d.; n=3). See also Sup. Fig. S6B. (**B**) Calculated dissociation constants for the indicated tubulins. Elp456 binds with similar affinity to recombinant *Drosophila* α1β1-tubulin, pig brain tubulin, mouse brain tubulin tubulin and mouse brain tubulin lacking K40 acetylation. However, the binding to tubulin purified from TTLL1/7-doubleKO mouse brains (lacking poly-glutamylation) and HeLa S3 cells (lacking poly-glutamylation, K40 acetylation and Δ2 deletion^49^’^50^) is similar (≥20 µM) and ∼18 times lower than other tubulins. Note that removal of the last tyrosine does not seem to have an effect in Elp456 binding (CPA treated tubulin purified from HeLa S3 cells, see methods). (**C**) Experimental assay to detect poly-glutamylation in polymerized microtubules *in vitro*. Stable, HiLyte488-labelled microtubule seeds are attached to a surface as in Fig. 1. Microtubules are then polymerized from the seeds and the levels of poly-glutamylation are measured using fluorescently labelled anti-polyglutamylated tubulin antibodies. See methods for details about the quantification and signal normalization. (**D**) Microtubules polymerized in the presence of Elongator are significantly enriched in poly-glutamylated tubulin when using a 50:50 pig brain tubulin:HeLa S3 tubulin mix. Correspondingly, when using HeLa S3 tubulin, which is not poly-glutamylated^49,50^, no effect is observed. P values for an Ordinary one-way ANOVA test followed by Turkey multiple comparison test are indicated.

To pinpoint the specific PTM controlling the interaction between tubulin and Elp456, we used tubulin purified from mouse brains lacking critical enzymes modifying tubulins (see methods). Remarkably, tubulin purified from TTLL1/7-double KO mouse brain, lacking poly-glutamylation^50^, displays a much weaker affinity towards Elp456 (*K*_d_ ≥ 25 μM, n=3. **Fig. 6A, B**) than the wild-type (1.8 ± 0.6 μM, n=3). However, using ATAT1-KO mouse brain tubulin, we showed that Elp456 binding is independent of *α*-tubulin acetylation (*K*_d_ = 1.7 ± 0.6μM, **Fig. 6A, B**). Similarly, Elp456 binding was independent of the tyrosination state of the *α*-tubulin C-terminal tail, since HeLa S3 tubulin and de-tyrosinated HeLa S3 tubulin (using carboxypeptidase A to remove the C-terminal tyrosine) display almost identical binding curves (*K*_d_ ≥ 22 μM, n=3. **Sup. Fig. S6B**). This suggests that the Elp456-tubulin interaction requires poly-glutamylated *αβ*-tubulin C-terminal tails (**Fig. 6B**).

Note that absence of most post-translational modifications in the C-terminal tail greatly reduces, but not completely abolishes, the interaction with Elp456 (**Fig. 6B, Sup. Fig. S6B**, HeLa S3 and HeLa S3+CPA). This suggests that Elp456 also recognizes the body of the tubulin, although weakly, which could explain the weak binding of Elp456 towards microtubules we observed in TIRF microscopy (**Sup. Fig. S4A, B**). These results suggest that the body of the tubulin molecule likely plays a role in the interaction, as otherwise full binding of Elp456 to microtubules would be expected if only the C-terminal tails participated in the interaction, independently of the polymerization state of tubulin.

### Elongator is a polyglutamylation-selective tubulin polymerase

Whilst Elp456 specifically binds to poly-glutamylated tubulin, the interaction of Elp123 with the microtubule lattice is independent of PTMs. Indeed, we observed that Elp123 is still able to bind to microtubules assembled from HeLa tubulin or pig brain tubulin partially digested with subtilisin to remove the C-terminal tail once polymerised (**Sup. Fig. S2E, F**). Therefore, whilst binding of Elp123 to microtubules is independent of their post-translational modifications, the binding of Elp456 is not. Given the molecular mode of action of Elongator (Figs 1-6), this surprising property could endow Elongator with the unique ability to selectively enrich microtubules with poly-glutamylated tubulin without harbouring a tubulin modifying enzymatic activity. In other words, by binding to microtubules irrespective of their PTMs, but selectively elongating them with poly-glutamylated tubulin subunits, Elongator could change the tubulin code of *dynamic* microtubules. This stands in contrast to the mode of action of the modifying enzymes such as members of the TTLL family, which preferentially modify long-lived, stable microtubules^51-53^.

To test this hypothesis, we established an assay to quantitatively measure the incorporation of poly-glutamylated tubulin in polymerized microtubules (**Fig. 6C**, see also methods). Briefly, microtubules were elongated from stable, HiLyte488-labelled GMP-CPP seeds in the presence or absence of Elongator, from a non-fluorescent source of heterodimers composed of a 50:50 mixture of HeLa S3 tubulin, which is weakly poly-glutamylated, and pig brain tubulin, which is heavily poly-glutamylated. After polymerization, the microtubules were stabilized with Taxol, incubated with fluorescently-labelled antibodies against poly-glutamylated tubulin and imaged by TIRFM. This allowed us to measure the amount of poly-glutamylation within the microtubules with and without Elongator (**Fig. 6C**). To normalize for sample-to-sample differences in fluorescence, the poly-glutamylated tubulin antibody signal was normalized against the HiLyte488 signal from the microtubule seeds (see methods). Note that we use a 50:50 HeLa S3/pig brain tubulin to decrease the ratio of poly-glutamylated tubulin in the initial pool and thereby facilitate the detection of any enrichment in the resulting microtubules.

Strikingly, we could detect a clear enrichment in poly-glutamylated tubulin in microtubules assembled in the presence of Elongator compared to the control (normalized intensity - 23.58 ± 4.93, n=627 versus 11.70 ± 2.13, n=618 for the control, **Fig. 6D, Sup. Fig. S6D**). Note that, as expected, microtubules grow longer and at higher density in the presence of Elongator (**Sup. Fig. S6D-E**), confirming that indeed the Elongator complex participated in the elongation of microtubules. A control condition with only HeLa S3 tubulin showed low poly-glutamylation-signal in both the absence (4.35 ± 1.93, n=638) and presence (4.08 ± 1.8, n=619) of Elongator (**Fig. 6D**), confirming the low abundance of poly-glutamylated tubulin in the sample. Crucially, microtubule density was similar in the presence or absence of Elongator after polymerization of HeLa S3 tubulin (**Sup. Fig. S6F-G**), which is in line with our findings that Elongator displays low affinity towards this tubulin (**Fig. 6B, Sup. Fig. S6B**). Note that the higher density of microtubules in HeLa S3 tubulin conditions reflects an intrinsically higher polymerization rate of HeLa S3 tubulin compared to pig brain tubulin, independent of Elongator (**Sup. Fig. S6F-G**). This confirms that Elongator does preferentially use poly-glutamylated tubulin as a substrate and, therefore, Elongator does not stabilize microtubules when the available tubulin pool is not poly-glutamylated. Altogether, these results highlight an unexpected property of the Elongator complex, namely that it functions as a microtubule polymerase selective for poly-glutamylated tubulin.

## Discussion

### Convergent evolution of the mode of action of microtubule polymerases

A key finding of our study is that Elongator directly binds to both the growing microtubule ends and to free *αβ*-tubulin heterodimers, and that the coupling between these two activities allows Elongator to modulate microtubule dynamics. Indeed, we found that: i) Elongator specifically recognises the growing ends of microtubules, both at the plus and at the minus ends (**Fig. 1C, E; Sup. Fig. S1B-C);** ii) the Elp123 sub-complex binds directly to microtubules (**Fig. 2, Sup. Fig. S2D, F**), but this binding is not sufficient to drive microtubule stabilization (**Fig. 2B, C);** iii) Elp456 binds to soluble *αβ*-tubulin heterodimers (**Fig. 3B, C; Sup. Fig. S3C, D**) but not microtubules; iv)when Elp123 binds to microtubules, it can recruit Elp456, which in turn brings tubulin (**Fig. 3A, Fig. 4A, Sup. Fig. S4);** v) both Elp123 and Elp456 are needed to stabilize microtubules (**Fig. 4B, D**); and vi) weakening the interaction between both sub-complexes results in a loss of microtubule stabilization (**Fig. 5**). This dual binding of Elongator to microtubule ends and *αβ*-tubulin heterodimers explains molecularly how Elongator enhances the growth speed and thereby the persistence of microtubules by increasing the local concentration of tubulin at both ends (**Fig. 5A**).

This mechanism of action relying on increasing the local density of free tubulin at microtubule tips is reminiscent of other microtubule polymerases, like CLASP and XMAP215. Like Elongator, CLASP (in the presence of EB1) and XMAP215 have indeed been shown to track microtubule ends and to recruit free tubulin once bound, thereby enhancing the incorporation of tubulin dimers to the growing ends of microtubules^6-10^. But there is a major difference, namely that Elongator does not contain any sequence motive or domains predicted to interact with *αβ*-tubulin heterodimers (i.e. TOG domains) or with end-binding proteins (i.e. SxIP motifs). It is striking that a complex devoid of TOG-domains like Elongator has evolved to converge towards a similar mode of action through a completely orthogonal way, in this case using two subcomplexes to achieve the same result as having two separate TOG-domains. This different mode of action might explain another key difference between Elongator and XMAP215, namely that while both increase the microtubule polymerisation rate, Elongator decreases the catastrophe rate, on the contrary to XMAP215, which increases it^54^, it is tempting to speculate that cells needed to evolve multiple microtubule stabilizers with different properties because despite the relatively high physiological concentrations of tubulin in cells, the frequency of microtubule catastrophe is still relatively high at these concentrations. Proteins like CLASP, XMAP215 and Elongator that locally increase the concentration of tubulin is probably a cost-efficient way to ensure robust microtubule elongation.

It is worth noting that we detected a higher effect on microtubule stability when using an excess of Elp456 over Elp123 to reconstitute Elongator (**Fig. 4B-D**). When purified using buffers containing physiological salt concentrations, Elongator usually comes as an asymmetric complex containing only one copy of the hexameric ring Elp456^26,28,29^(**Fig. 1A**). However, it is clear from the structure that an extra surface for Elp456 binding is available on the other lobe of the Elp123 dimer. Indeed, it has been reported that an extra Elp456 ring could be observed in some 2D classes averages obtained by negative staining when an excess of Elp456 was used and/or the complex purified in low-salt conditions^29^. Since the microtubule dynamic experiments in this study were conducted in a low salt buffer which is critical for microtubule polymerization, it is tempting to suggest that a second Elp456 sub-complex could be recruited to Elp123 when Elp456 is added in excess. Indeed, this would further increase the local concentration of *αβ*-tubulin heterodimers at the growing ends of microtubules, enhancing the effect on microtubule stabilization that we observed (**Fig. 4B-D**). Further structural characterization is necessary to determine if a complex consisting of an Elp123 dimer, two copies of Elp456, and two copies of *αβ*-tubulin is sterically possible and can be used to assemble microtubules.

### An orthogonal way of writing the tubulin code

Whilst characterizing the mechanism by which Elongator binds to microtubules and tubulin heterodimers, we found a very unexpected behaviour, namely that Elongator can discriminate between tubulin heterodimers carrying different PTMs. On one hand, the Elp456 subcomplex displays ∼18-fold reduced affinity towards tubulin lacking poly-glutamylation, while changes in acetylation and tyrosination had little effects (**Fig. 6A, B, Sup. Fig. S6B**). On the other hand, the Elp123 subcomplex binds to polymerized microtubules independently of their PTM status (**Sup. Fig. S2E, F**), This made the intriguing prediction that microtubules polymerized by Elongator should become selectively enriched in polyglutamylated tubulin, which we could confirm experimentally (**Fig. 6C, D**).

Overall, the Elongator complex showed an unprecedented behaviour for both tubulin polymerases and tubulin PTM regulators. Indeed, it is well established that tubulin polyglutamylases, which are members of the tubulin-tyrosine ligase-like (TTLL) family, prefer modifying polymerized microtubules over free tubulin^21,55^. Hence, the current paradigm postulates that microtubules can only get modified *after* their assembly from a homogenous pool of free tubulin. To achieve spatial modification of microtubules with this paradigm, TTLL proteins must be targeted to the desired microtubules, for instance by other microtubule-associated proteins. This has been demonstrated for CEP41 (*Centrosomal Protein of 41 kDa*), CCSAP (*Cilia and Spindle-Associated Protein*) and PGs1 (*Tubulin Polyglutamylase complex subunit 1*) directly recruiting and/or activating TTLL proteins to specific structures in the cells^56-58^.

Our discovery that Elongator assembles dynamic microtubules specifically from polyglutamated tubulin now shifts this paradigm and offers an orthogonal way to modify microtubules. Indeed, since it can bind to microtubules regardless of their PTM composition, and that its activity biases microtubule towards polyglutamylated tubulin, Elongator effectively changes the PTM composition of microtubules as they are being polymerized. In other words, whilst most tubulin-modifying enzymes change the tubulin code of *stable* microtubules, Elongator changes the code of *dynamic* ones. This could be important for very dynamic, short-lived microtubule structures, such as the mitotic spindle. Indeed, interpolar microtubules at the mitotic spindle are enriched in poly-glutamylated tubulin^14,59^. Since Elongator is recruited to the mitotic spindle^30^ this opens the possibility for Elongator and other factors acting in a similar way to actively remodel the PTM landscape of the newly synthesized interpolar and astral microtubules, a crucial element for the accurate completion of cell division^12,60-62^. In addition, we previously reported that Elongator induces cytoskeleton symmetry breaking during asymmetric cell division, suggesting that Elongator activity can be modulated by cortical polarity cues^30^. It is thus tempting to speculate that Elongator could imprint polarity onto the landscape of PTMs of the microtubule network downstream of polarity cues. This could explain the markedly nonhomogeneous landscape of microtubule PTMs in polarized cells such as neurons^12,63^.

## Materials and Methods

### Plasmids

For baculoviral expression of Elp123, genes encoding for Elp1 (CG10535), Elp2 (CG11887) and Elp3 (CG15433) were codon optimized for *Drosophila melanogoster*, synthetized by Twist Bioscience (San Francisco, CA)and sub-cloned into pACEBacl vectors either untagged (Elp1 and Elp2) or containing a His-PC-SNAP N-terminal tag (Elp3). The final pBiglA vector containing the three subunits (referred here as Elp123) was assembled using an enhanced Gibson Assembly mix^37,64^. Empty pbiG plasmids were gifts from Dr Andrew Carter, MRC-LMB, Cambridge, UK. The Elp123 pBig1A vector was then transformed into DH10EmBacY cells (Geneva Biotech) and plated onto agar plates containing 50 μg/ml kanamycin, 10 μg/ml tetracycline. 7 μg/ml gentamycin, 40 μg/ml isopropyl *β*-d-1 -thiogalactopyranoside (IPTG) and 100 μg/ml Blue-Gal. White colonies, in which the vector has been integrated with the baculovirus genome, were grown overnight in 4 ml of LB supplemented with 50 μg/ml kanamycin. 10 μg/ml tetracycline, 7 μg/ml gentamycin. Bacmid DNA was prepared using QIAprep Spin Miniprep Kit (27106, Qiagen) buffers according to the MultiBac protocol.

The plasmid used for Elongator purification in D.mel-2 cells was previously described in ref.^30^.

For co-expression of Elp4 (CG6907), Elp5 (CG2034) and Elp6 (CG9829) in bacteria, a modified pGEX vector was used as previously described^30^. Briefly, Elp4 was first cloned into a modified pGEX vector containing an N-terminal GST-tag followed by a TEV protease cleavage sequence. Then a synthetic fragment comprising both Elp5 and Elp6 ORFs, each flanked by a Ribosome Binding Sequences (RBS) and a stop codon, was cloned in 3’ of Elp4. This generates a single transcription unit (Promoter-GST-TEV-Elp4-STOP-RBS-Elp5-STOP-RBS-Elp6-STOP-Terminator) expressing all three subunits. This vector is referred to as pGEX-Elp456 in this study. This strategy was adapted to express all variants of Elp456 used in this study, including mScarlet-Elp456^30^, eGFP-Elp456, as well as Elp456 *ΔpolyN* and Elp456 *solo* mutants. For Elp456 *ΔpolyN*, the ΔpolyN region was deleted using divergent PCR using the pGEX-Elp456 vector as a template and primers that flank the deleted region in Elp4 (forward: 5’-GGGGGAACTCGATTTACTA-AATTC-3’, reverse: 5’-ACAGGCTCCCAGGATAGC-3’). The *solo* versions of Elp4 (K364E, R397E & E410R), Elp5 (K151E, K179E and T181A)and Elp6(K199E, S200W and K228E)were synthetized and cloned using the same strategy as for the wild-type.

For recombinant *Drosophila* tubulin expression and purification, a codon-optimised gene coding for tubulin α1 84B (Gen-Bank entry NM_057424) carrying a tandem N-terminal His_6_-tag and a Protein C epitope tag (PC tag, EDQVDPRLIDGKG) was custom-synthesised (Twist Bioscience) and cloned into a pMT Puro vector^30^.

### Insect cell transfection

For baculoviral expression of Elp123, Sf9 cells were seeded at 5·10^5^ cells/ml in a 6-well plate in a total volume of 2 ml of Sf-900-II SFM media (10902088, ThermoFisher Scientific). Bacmids (see above) were transfected using FuGENE^®^ HD using manufacturer’s protocol (E2311, Promega). After 9 days, the supernatant was recovered and used to infect a 50 ml culture of Sf9 cells at 2·10^6^ cells/ml. After 72h, the virus was harvested by pelleting the cells at 300g for 10 minutes. This virus was stored at 4°C in the dark and used at a 1:100 dilution for large-scale cell infection for protein production.

For holo-Elongator production in insect cells, we followed the protocol we previously established^50^. Briefly, D.mel-2 cells (CRL-1963, ATCC) were grown at 25°C in Insect-Xpress medium (181562, Lonza) supplemented with 1% Pen/Strep (15140122, Gibco) and 0.25 μg/ml Amphotericin B (15290018, Gibco). Cells were transfected with a pMT Puro His-PC-SNAP-Elp3 vector using ExpiFectamine™ Sf(A389l5, Gibco) using manufacturer’s instruction, followed by selection in 5 μg/ml Puromycin (A1113803, ThermoFisher).

### Mouse lines

Animal care and use for this study were performed in accordance with the recommendations of the European Community (2010/63/UE) for the care and use of laboratory animals.

Experimental procedures were specifically approved by the ethics committee of the Institut Curie CEEA-IC #118 (authorisation no. 04395.03 given by National Authority) in compliance with the international guidelines. Adult males and females (2-8 months) were used in this study. All mouse lines used in this study (wild-type, *Ttll1*^*−/−*^ *Ttll7*^*−/−*^and *Alat1*^*−/−*^) were described before^65,66^.

### Protein purification

For purification of Elp123, Sf9 cells co-expressing Elp1, Elp2 and His-PC-SNAP-Elp3 were pelleted, resuspended in lysis buffer (0.05 M K-HEPES, 0.1 M K-Acetate, 0.002 M MgCl_2_, 0.01 M CaCl_2_ 5%glycerol, 120μg/ml Benzamidine, 20μg/ml Chymostatin, 20 μg/ml Antipain, 0.5 μg/ml Leupeptin, 240 μg/ml Pefabloc and 1 mM PMSF, pH 8.0). Cells were lysed by extrusion using a Teflon Dounce homogenizer and clarified by centrifugation at 30000g for 40 minutes at 4°C using a Ja 25.50 rotor (Beckman). The clarified lysate was incubated with 2 ml of pre-equilibrated Protein C affinity resin (Roche) for 2h at 4°C. Then, the resin was packed in an empty column (Bio-rad), washed with 20 ml of wash buffer (0.05 M K-HEPES, 0.1 M K-Acetate, 0.002 M MgCl_2_, pH 8.0) supplemented with 0.001 M CaCl_2_, followed by 50 ml of wash buffer without CaCl_2_. The Elp123 sub-complex was then eluted with elution buffer (wash buffer supplemented with 0.01 M EGTA pH 8.0) and analysed by SDS-PAGE. The Elp123 containing fractions were further purified using a Heparin column (HiTrap Heparin HP, Cytiva) equilibrated in Elution buffer and eluted using a linear gradient between Heparin A (0.05 M K-HEPES, 0.1 M K-Acetate, 0.002 M MgCl_2_, 0.01 M EGTA, pH 8.0) and Heparin B (0.05 M K-HEPES. 1 M K-Acetate, 0.002 M MgCl_2_, 0.01 M EGTA, pH 8.0) buffers. The elution was analysed by SDS-PAGE, and the fractions of interest were pooled and injected in Superose 6 10/300 column (Cytiva) equilibrated and eluted in Elongator buffer (0.02 M K-HEPES, 0.15 M K-Acetate, 0.001 M DTT, pH 8.0). The peak fractions were collected, concentrated (Amicon Ultra-4 30kD MWCO, Millipore) and labelled using SNAP-Surface AlexaFluor 488 (NEB) using manufacturer’s instructions. The labelled protein was then desalted into Elongator buffer to remove non-bound dye, concentrated, flash-frozen in liquid N_2_ and stored in small aliquots at -80°C.

Holo-Elongator was purified from D.mel-2 cells transfected with His-PC-SNAP-Elp3 using a previously established protocol^30^. Briefly, the transfected D.mel-2 cells were grown to litre-scale over several weeks, then the expression was induced by the addition of 0.6 mM CuSO4. After 4 days, the cells were pelleted, lysed, and purified using a PC-resin as described for Elp123 except that all buffers contain 20% glycerol. After PC-resin elution, Elongator-containing fractions were concentrated and further purified by sucrose gradient by layering the protein on top of a manually prepared 10-30% discontinuous sucrose gradient. After ultracentrifugation for 1h at 258,488 x g at 4°C using a TLS-55 rotor (Beckman), the fractions were analysed by SDS-PAGE and the positive fractions were concentrated, labelled with SNAP-Surface Alexa Fluor 488, and desalted using the same procedure described for Elp123.

All variants of the Elp456 sub-complex (wild type, mScarlet tagged, eGFP tagged, ΔpolyN and *solo* mutant) were purified using the same protocol. BL21 (DE3) Rosetta2 (Invitrogen) *Escherichia coli* were transformed with the respective vector and grown at 37°C in 2xYT medium to OD_600_=0.8. Protein expression was induced by the addition of 0.5 mM IPTG at 20°C for at least 16 hours. The cells were collected by centrifugation, resuspended in lysis buffer (0.05 M K-HEPES pH 7.5.0.1 M K-Acetate. 5% glycerol. 0.001 M DTT. 0.01 M MgCl_2_, 1% Triton X-100 and cOmplete™ Protease Inhibitor Cocktail Tablets (Roche) and lysed by sonication. The lysate was then clarified (using a JA 25.50 Beckman rotor at 40000x g for 30 min, 4°C) and the soluble fraction was incubated with 2 ml of Glutathione Sepharose 4B resin for 2h at 4°C under constant agitation. The resin was washed with 4 times 20 ml of wash buffer (0.05 M K-HEPES pH 7.5, 0.1 M K-Acetate, 5% glycerol, 0.001 M DTT, 0.01 M MgCl_2_) and the protein was eluted by incubating the resin with rTEV protease (overnight, 4°C) to remove the N-terminal GST-tag in Elp4 (releasing the protein from the resin). The next morning, the eluted protein was recovered and further purified by size-exclusion chromatography on a Superdex 200 16/60 column (Cytiva) in Elongator buffer. Finally, the peak fractions containing the three proteins (Elp4, Elp5 and Elp6 at seemingly equimolar quantities) were concentrated, flash-frozen in liquid N_2_ and stored in small aliquots at -80°C.

Unlabelled porcine tubulin, AMCA-, rhodamine- and Hi-Lyte647-labelled porcine tubulin were purchased from Cytoskeleton (T240, TL440M, TL590M and TL670M, respectively). All tubulins were reconstituted at 5 mg/ml in BRB80 buffer (80 mM K-Pipes, 1 mM MgC12,0.5mM EGTA, pH 6.9) supplemented with either 1 mM GTP (Roche), 1 mM GMPPCP (NU-405S, Jena Bioscience) or 1 mM BeF_4_ (prepared by mixing 50 mM BeSO_4_ and NaF 100 mM to make 10 mM BeF_3_), flash frozen and kept in liquid N_2_. Microtubules seeds stocks were prepared at 5 mg/ml tubulin concentration (20% fluorescent-tubulin, 20% biotinylated tubulin) in BRB80 buffer supplemented with either 1 mM GMPCPP or 1 mM BeF_4_, aliquoted in 1 μl aliquots and stored in liquid N_2_.To polymerize the seeds, one aliquot of microtubule seed stock is incubated at 37°C for 30 minutes, sedimented on a table-top centrifuge by centrifugation (8 minutes at 14000g), and resuspended in BRB80 buffer supplemented with 1 mM nucleotide.

For recombinant *Drosophila* tubulin purification, D.mel-2 cells were transfected and grown using the same strategy as for His-PC-SNAP-Elp3. To purify the *αβ*-tubulin heterodimers, the cells were lysed in lysis buffer (80 mM K-PIPES pH 6.9. 10 mM CaCl_2_, 10 μM Na-GTP, 10 μM MgCl_2_, 0.12 mg/mL benzamidine, 20 μg/mL chymostatin, 20 μg/mL antipain. 0.5 μg/mL leupeptin, 0.24 mM Pefabloc SC, 0.5 mM PMSF) and lysed by using a Dounce homogeniser. The lysate was rocked for 1h at 4°C to ensure complete microtubule depolymerisation, then clarified by centrifugation at 66,000 x g for 30 min using a JA 25.50 rotor (Beckman). The supernatant was incubated with 2 mL of pre-equilibrated Protein C affinity resin (Roche) for 3h at 4°C. After incubation, the resin was packed into an empty column (Bio-Rad), washed with 50 ml of tubulin wash buffer (80 mM K-PIPES pH 6.9,10 μM Na-GTP, 10 μM MgCl_2_, 1 mM CaCl_2_), 50 mL of tubulin ATP-buffer (wash buffer + 10 mM MgCl_2_ and 10 mM Na-ATP), 50 mL of low-salt buffer (wash buffer + 50 mM KCI), 50 ml of high-salt buffer (wash buffer + 300 mM KC1), 50 mL of Tween buffer (wash buffer + 0.1% Tween-20 and 10% glycerol) and finally 50 mL of tubulin wash buffer without CaCl_2_. Tubulin was eluted with tubulin buffer (80 mM K-PIPES, 10 μM Na-GTP. 10 μM MgCl_2_, 5 mM EGTA, pH 6.9). The protein-containing fractions were pooled and further purified on a Superdex 200 column (GE Healthcare), equilibrated, and eluted in tubulin buffer. The peak fractions were pooled and concentrated to 10 mg/ml, flash-frozen in liquid N_2_ and stored in liquid N_2_. The resulting *αβ*-lubulin heterodimers are formed by isotype-pure tubulin α1-84B (Uniprot TBA1_DROME) and the co-purified β tubulin (tubulin β1-56D, Uniprot TBB1_DROME^67^. We previously characterised this tubulin by CryoEM which shows that purified recombinant tubulin purified this way is functional and assembles in 13-protofilament microtubules in the presence of GTP^67^.

### Tubulin purification from murine brains

Tubulin was purified from mouse brains (see Mouse lines) via cycles of temperature-dependent microtubule polymerisation and depolymerisation as previously described^50,68^. Briefly, the animals were sacrificed and the brains extracted immediately and added in ice-cold lysis buffer (BRB80 (80 mM K-PIPES pH 6.8, 1 mM K-EGTA, 1 mM MgCl2), 1 mM *β*-mercaptoethanol. 1 mM PMSF and 1× protease inhibitor cocktail: 20 μg/ml leupeptin, 20 μg/mI aprotinin and 20 μg/mI 4-(2aminoethyl)-benzenesulfonyl fluoride; Sigma-Aldrich) in a ratio of 2 ml buffer per 1 g of tissue. The brains were homogenised using an Ultra-Turrax® blender and the lysates cleared by centrifugation at 112,000 ×g in TLA-55 fixed-angle rotor (Beckman Coulter) for 30 min at 4°C (i.e. cold centrifugation). To this first supernatant (SN1) final concentration of 1 mM GTP and 1/3 volume pre-warmed 100% glycerol were added, mixed and incubated for 20 min at 30°C to allow microtubule polymerisation. Polymerised microtubules were pelleted for 30 min at 112,000 × g, 30°C, (i.e, warm centrifugation) and the supernatant SN2 discarded. The microtubule-containing pellet was resuspended in cold BRB80 (depolymerisation 1; 1/10 of the initial SN1-volume)and incubated on ice with occasional pipetting up-and-down over 20 min to mechanically assist microtubule depolymerization. Solubilised tubulin was cleared via cold centrifugation, the supernatant SN3 was transferred to a fresh tube and adjusted to final concentrations of 1 mM GTP and 0.5 M PIPES, complemented with 1/3 volume pre-heated 100% glycerol. The microtubule polymerization step was repeated by incubation of the solution for 20 min at 30°C and the resulting microtubules were pelleted via warm centrifugation. Note that the microtubules yielded in this step by polymerisation in presence of the high molarity buffer are largely free of associated proteins. The supernatant SN4 containing these associated proteins was discarded and the pellet was resuspended in cold BRB80 (depolymerisation 2; 1/40 of the initial SN1-volume), depolymerised on ice for 20 min as before, and cleared via cold centrifugation. The resulting SN5 was adjusted to 1 mM GTP, supplemented with 1/3 volume of pre-heated 100% glycerol, and incubated for 20 min at 30°C, after which the microtubules were pelleted by warm centrifugation and resuspended in ice-cold BRB80 (depolymerisation 3; 1/ 40 of the initial SN1-volume). After 15 min on ice, the soluble tubulin was cleared by a final cold centrifugation and the tubulin yield was estimated with a NanoDrop ND-1000 spectrophotometer (Thermo Scientific; absorbance at 280 nm; MW = 110 kDa; ε = 115,000 M − 1/cm − 1). Samples were aliquoted, snap-frozen in liquid N2 and stored at −80°C.

### Purification of tyrosinated and de-tyrosinated tubulin from Hela cells

Tubulin was purified from Hela S3 cells using cycles of polymerisation and depolymerisation similar to the approach used for brain tubulin^50^. Briefly, Hela S3 cells were grown in suspension in 1-L spinner bottles for 7 days, collected, and the cell pellet (approx. 10 ml for 4 spinner bottles) was lysed in the same volume of ice-cold lysis buffer (same composition as for the brain tubulin prep, see above) using a French press. The subsequent steps were the same as described above for brain tubulin, except that during the depolymerisation step 1, the microtubule pellet was resuspended in 1/60 of the SN1 volume, for depolymerisation 2 in 1/100 of the SN1 volume, and for depolymerisation 3 in 1/300 of SN1 volume. Tubulin purified from Hela cells is highly tyrosinated^49^. To obtain detyrosinated tubulin, the solubilized tubulin in SN3 was incubated with 1/300 volume of Carboxypeptidase A (Sigma C9268) at 30°C for 5 min before the incubation in high molarity PIPES buffer, and the rest of the purification procedure was followed as described above.

### SDS-PAGE and Western blot

SDS-PAGE was performed using NuPAGE 4-12% Bis-Tris gels (Life Technologies) according to the manufacturer’s instructions. Instant Blue (Sigma) was used for total protein staining of gels.

For Western Blot, gels were transferred on nitrocellulose membranes using iBLOT (Life Technologies). The membranes were first stained with Ponceau (Sigma) to assess the quality of the transfer, then washed in TBS and blocked in TBS supplemented with 5% semi-skimmed milk for 30 min at RT. For fluorescent western blot with Atto-647N-labelled anti-α K4O acetylated tubulin antibodies^69^, membranes were incubated overnight at 4°C with 1 μg/ml antibody in TBS supplemented with 0.2 % BSA. The membranes were then imaged using a Typhoon scanner.

### Microtubule co-sedimentation assay

All proteins were pre-clarified at 20000 x g at 4°C for 10 minutes before use in biochemical assays. Reconstituted porcine tubulin or HeLa tubulin was diluted to 50 μM in BRB80 buffer supplemented with 1 mM GTP. To initiate microtubule polymerization, polymerization buffer (BRB80 supplemented with 60% glycerol) was added to a final concentration of 5% glycerol. After 20 minutes incubation at 37”C in a water bath, the microtubules samples were diluted to 10 μM in BRB80 buffer supplemented with 40 μM Taxol.

To prepare subtilisin-digested porcine tubulin, freshly prepared Taxol-stabilized microtubules were incubated with 0.05 mg/ml subtilisin (P538, Sigma) for 40 minutes at 37°C in a water bath. The reaction was stopped by the addition of 2 mM PMSF (93482, Sigma).

At this point, all microtubule samples were cleared by spinning for 15 minutes at 20000 x g at Room Temperature to remove non - polymerized tubulin. The supernatant was then discarded and the microtubules resuspended in BRB80 buffer supplemented with 40 μM Taxol.

To test the binding of Elp123 to microtubules, 50 μL solutions were prepared with 5 μM microtubules and 160 nM Elp123, and the samples were incubated at Room Temperature for 30 minutes. After incubation, the samples were transferred into 20 mm polycarbonate ultracentrifuge tubes (Beckman) pre-filled with 150 μl of cushion solution (BRB80 supplemented with 60% glycerol and 40 μM taxol). The samples were centrifuged at 100.000 x g for 30 minutes at 25°C using a TLA100 rotor. After centrifugation, the top 20 μl were recovered and saved for SDS-PAGE/Western blot analysis, designated as “supernatant”. The remaining 10 μl atop the glycerol cushion were removed and discarded, and the interfaces were washed three times with 100 μl of BRB80 buffer. The glycerol cushion solutions were then removed and the pellets gently washed with 3×100 μl of BRB80 buffer. Finally, the pellets were resuspended in 50 μl of BRB80 buffer, saved and designated as “pellet” for further analysis.

### Microtubule dynamics assays

All proteins were pre-clarifled at 20000 x g at 4°C for 10 minutes before use in microscopy assays. Glass 22×22 mm coverslips (NEXTERION, Schott) were incubated at least for 48h at room temperature with gentle agitation in a 1:10 (w/w) mix of mPEG-Silane (30 kDa, PSB-2014, Creative PEGWorks) and PEG-Silane-Biotin (3.4 kDa, Laysan Bio) at a final concentration of 1 mg/ml in 96 % (v/v) ethanol and 0.2 % (v/v) HC1. The day of the experiment, the coverslips were washed with ethanol and ultrapure water, dried with a nitrogen gas gun, and assembled into an array of flow cells on mPEG-Silane passivated slides using double-sided tape (Adhesive Research AR-90880 precisely cut with a Graphtec CE6000 cutting plotter). The chamber was first perfused with BRB80 buffer, and then washed with 5% Pluronic-F127 and BRB80. Neutravidin (25 μg/ml) was then added to the chamber and incubated for 5 minutes, then washed out with 10 chamber volumes of BRB80. Biotin-stabilized microtubule seeds (20% with rhodamine-tubulin, 20% biotin-tubulin) were injected in imaging buffer (BRB8O enriched with 0.1 mg/ml K-Casein, 0.1 mg/ml BSA, 40 μM DTT, 64 mM D-glucose, 160 μg/ml glucose oxidase, 20μg/ml catalase and 0.2% methylcellulose) and let for 5 minutes to bind to the neutravidin, followed by a final wash with imaging buffer. Dynamic microtubules were elongated from the seeds by injecting in the chamber a solution of 10% labelled HiLyte647-GTP Tubulin and different Elongator sub-complexes (or an equivalent volume of Elongator buffer as a control) and microtubule dynamics were monitored by TIRFM. Note that an aliquot of the last buffer exchange step during the purification of each protein was always kept to serve as a true negative control in all tubulin dynamics experiments. The microscope stage was kept at 37°C with an objective heater in addition to the heating chamber containing the microscope.

### Microtubule-decoration experiments

To further increase the signal-to-noise ratio in our TIRFM images for microtubule-decoration experiments (Fig. 1E, Fig. 2A, Fig. 3A, Fig. 4A and Sup. Fig S4) we modified the passivation of the glass coverslips as follows: glass coverslips were initially incubated with a 1 mg/ml solution of mPEG-Silane. After assembly, the chamber was first perfused with BRB80, then incubated with a solution of 0.1 mg/ml PLL-g-PEG (Susos) to further passivate the surface. After three minutes, the chamber was washed with 10 volumes of BRB80 then passivated with 1 mg/ml K-Casein. After three more minutes, GMPCPP-stabilized microtubule seeds diluted in imaging buffer were introduced in the chamber and left for 5 minutes to settle. Then, the chamber was washed with imaging buffer and the proteins of interested were sequentially added to the chamber. Concentrations used are: Fig. 2A, 100 nM 488-SNAP-Elp123; Fig. 3A, 100 nM 488-SNAP-Elongator, 15 μM 647-GTP-Tubulin; Fig. 4A and Sup. Fig S4, 100 nM 488-SNAP-Elp123, 500 nM mScarlet-Elp456, 15 μM 647-GTP-Tubulin.

### Microtubule poly-glutamylation detection experiments

All proteins were centrifuged for 10 minutes at 20000 xg at 4°C before use in biochemical assays. Glass coverslips and experimental chambers were prepared following the same protocol as for microtubule dynamics experiments. After addition of biotin-stabilized microtubule seeds (20% with HiLyte488-tubulin, 20% biotin-tubulin), the chamber was washed with imaging buffer and then 15 μM of tubulin (either 50:50 pig brain tubulin:HeLa S3 tubulin, or 100% HeLa S3 tubulin) together with either 50 nM Elongator complex or Elongator buffer were injected in the chamber and incubated at 37°C. After 10 minutes, a mix containing 50 μM taxol and 30 nM anti poly glutamylated alpha tubulin (GT335, AdipoGen Life Sciences) labelled with ATTO565 in imaging buffer was injected, and after 10 minutes, the chamber was finally washed with imaging buffer containing 50 μM taxol. The microscope stage was kept at 37°C.

To quantify the poly-glutamylation state of polymerized microtubules, we measured the intensity of hundreds of microtubules per condition (from two or three independent experiments) in both the ATTO565-fluorescence (polyglutamylated tubulin) and HiLyte488 (microtubule seeds) fluorescence channels. For this, we traced a 2 pixel-wide line along the length of each microtubule and measured the underlying signal using the linescan function in Fiji^70^. For each microtubule, the 565-fluorescence intensity was averaged along the length of the microtubule and the background signal value was subtracted (background was measured at the edges of the linescan devoid of microtubule). Then, the calculated signal of all microtubules in a field of view (FOV) was averaged, and the result was normalized against the average 488-fluorescence Intensity from the microtubule seeds of that FOV, generating an averaged value per FOV. For each condition, we then averaged this mean normalized 565-fluorescence microtubule intensity across multiple FOVs (n=6-9) and compared them (non-transparent dots Fig.6D, individual values for each microtubule are also displayed as transparent dots). Note that analysing this dataset by a two-way ANOVA confirmed that the vast majority of the variation comes from comparing different conditions (86.25%), and not from different FOVs from the same condition (0.21%), whose differences are not statistically significant (i.e. FOV1 from Control vs FOV2 from Control, p_value = 0.9945). In addition, note also that the Hi-Lyte488 fluorescence signal from the seeds is very homogeneous between FOVs and samples, highlighting the homogeneity of the samples, microscope conditions and TIRF field (Control: 178.62 ± 11.19 a.u from 354 seeds, Elongator: 174.42 ± 13.49 a.u from 375 seeds, HeLa S3 control: 175.31 ± 17.91 a.u from 281 seeds, and HeLa S3 Elongator: 179.84 ± 10.86 a.u from 288 seeds). HiLyte488 fluorescence differences are not significant between FOVs of the same conditions, and between all conditions (Ordinary one-way ANOVA followed by a Turkey test for multiple comparisons). Number of analysed microtubules, FOV and independent experiments are: Control (N=3 independent experiments, n=9 FOV, n=618 microtubules), Elongator (N=3 independent experiments, n=9 FOV, n=627 microtubules), HeLa S3 Control (N=2 independent experiments, n=6 FOV, n=638 microtubules) and HeLa S3 Elongator (N=2 independent experiments, n=6 FOV, n=619 microtubules).

To quantify the total poly-Glutamylated tubulin signal in a field of view (Sup. Fig. S6C-F), the total signal from the antibody channel was measured in Fiji^70^ and the background was subtracted (background signal was extrapolated from the mean average per pixel of the signal from a region of interested devoid of signal multiplied by the total amount of pixels in the field of view). The resulting number was divided by the amount of microtubule seeds observed.

### Tubulin binding experiments

The binding of Elp456 to tubulin was first assessed using gel filtration experiments. Pig brain tubulin was diluted to 50 μM in ice-cold BRB80 and kept on ice for 30 minutes to favour microtubule depolymerization. Then, Elp456 was added to create a final solution of 30 μM Elp456 and 35 μM tubulin. The proteins were incubated on ice for 30 minutes, then injected in a superpose 6 increase 3.2/300 column equilibrated and eluted in binding buffer (40 mM K-Pipes, 1 mM MgCl2,1 mM EGTA, 20 μM GTP, 20 μM MgCl_2_, pH 6.9)(Fig.3B). The elution profile was analysed by SDS-PAGE (Fig.3C).

The binding of Elp456 and Elp123 to tubulin was also analysed by protein immunoprecipitation (Fig.S3). For Elp123, 30 μM pig brain tubulin was incubated alone or in the presence of 1 μM PC-Elp123 in binding buffer supplemented with 1 mM CaCl_2_ (required for PC-binding to the epitope) for 30 minutes on ice. Then, the solution was diluted 10-fold in binding buffer and 20 μL of PC-resin (pre-equilibrated in binding buffer) was added to the mixture. The proteins were left to interact with each other and bind to the resin for 2h in constant rotation at 4°C. Then, the sample was centrifuged for 10 minutes at 20000 x g at 4°C, the resin was washed thrice with binding buffer (CaCl_2_ was removed in the last wash step to facilitate elution) and finally the PC-bound proteins were released by incubating the resin with binding buffer supplemented with 5 mM EGTA for 30 minutes (twice). The results were analysed by SDS-PAGE, 488-fluorescenee (using a typhoon scanner) and western-blot anti αTubulin. To assess the binding of Elp456 to tubulin, the same protocol was followed except that non-tagged Elp456 and recombinant *Drosophila* PC-Tubulin (both at 30 μM) were used.

### Microscale thermophoresis

To measure the Elp456 binding affinity to different tubulins or to Elp123, microscale thermophoresis (MST) assays were performed on a Monolith X (Nanotemper Technologies) using standard capillaries (MO-K022) at 25°C. 16 or 24 two-fold dilutions series of tubulin or Elp123 were performed in binding buffer supplemented with 0.05% (v/v) Tween-20, eGFP-Elp456 (WT or mutated) was used at a constant concentration of 50 nM. Thermophoresis was measured using automatic LED power and irradiating the capillaries for 20 seconds at medium laser power (50%). The dissociation constant values were obtained by analysing the change in thermophoresis using the MO.Affinity (v3.0.5) software provided by the manufacturer.

### Total internal reflection fluorescence microscopy (TIRFM) and image processing

All imaging was performed using a custom TIRF microscope composed of a Nikon Ti stand equipped with perfect focus, a fast piezo z-stage (ASI) and a Plan Apochromat lambda 100X NA 1.45 objective. TIRF illumination was achieved with a azimuthal TIRF illuminator (iLas2, Roper France) modified to have an extended field of view to match the size of the camera (Cairn). Images were recorded with a Photometries Prime 95B back-illuminated sCMOS camera run in pseudo-global shutter mode and synchronized with the rotation of the azimuthal illumination. TIRF angle was set independently for all channels so that the depth of the TIRF field was identical for all channels. Excitation was performed using 405 nm (100 mW OBIS LX). 488 nm (150 mW OBIS LX). 561 nm (100 mW OBIS LS) and 637 nm (140 mW OBIS LX) lasers fibered within a Cairn laser launch. To minimize bleedthrough, single band emission filters were used (Chroma 525/50 for Alexa 488/GFP/Alexa514; Chroma 595/50 for mScarlet/rhodamine/mRFP/mCherry and Chroma 655LP for Hi-Lyte647/Alexa647/ATTO647N) and acquisition of each channel was performed sequentially using a fast filter wheel (Cairn Optospin). To enable fast acquisition, the entire setup is synchronized at the hardware level by a Zynq-7020 Field Programmable Gate Array (FPGA) stand-alone card (National Instrument sbrio9637) running custom code. Sample temperature was maintained at 25°C (or 37°C for microtubule dynamics) using a heating enclosure (MicroscopeHeaters.com, Brighton, UK). Acquisition was controlled by Metamorph software (v7.10.1.161).

Dual IRM (Interference reflection microscopy) and 488-TIRF imaging were performed in a LUMICKS C-Trap Edge setup. The chambers were prepared using the same protocol as for TIRFM. For IRM, 10 frames of 100 ms were collected and averaged, and the background was subtracted using LUMICKS BlueLake software. Sample temperature was maintained at 25°C.

### TIRM image processing

Images were processed using Fiji^70^ (ImageJ version: 1.53f) and Matlab 2020b (Mathworks). Figures were assembled in Adobe Illustrator 2021. All lookup tables applied to images in this paper come from the collection from James Manton (https://github.com/jdmanton/imageJ_LUTs).

Spatial drift during acquisition for TIRF movies was corrected using a custom GPU-accelerated registration code based on cross correlation between successive frames. Drift was measured on one channel and applied to all the channels in multichannel acquisitions. Code is available on our github page (https://github.com/deriverylab/GPU_registration).

### Cryo-Electron microscopy grid preparation, data collection and analysis

Elp456 samples (WT or mutated) were diluted to 1 mg/ml in Elongator buffer (0.02M K-HEPES, 0.15M K-Acetate, 0.001M DTT, pH 8.0). Samples were applied to freshly glow-discharged (Edwards S150B, 45 s, 30−50 mA, 1.2 kV, 10−2 mbar) copper holey carbon grids (Quantifoil, Cu-R1.2/1.3) under 100% humidity at 4°C.

Excess sample was blotted away, and the grids were subsequently plunge-frozen in liquid ethane using a Vitrobot Mark III (Thermo Fisher Scientific). Note that different strategies were tried to reduce the preferential orientation problem observed in the sample (including, but not limited to changing buffer composition and pH; detergents; protein concentration, tags, and batches; grid type; grid support; freezing conditions….). Elp456 WT and *solo* mutant samples (Fig. S5D) were imaged on a ThermoFisher Scientific Glacios electron microscope operating at 200 keV with a calibrated magnification of 92000x, corresponding to a pixel size of 1.58 Å, using a Falcon III direct electron in linear mode with a dose rate of 25.35 e/Å^2^/s and defocus ranging from -1 to -4 μm. Total exposure time was 2 seconds, resulting in an accumulated dose of ∼50 e/Å^2^. EPU software (ThermoFischer Scientific) was used for automated data acquisition. 311 or 337 images were collected for the WT and *solo* versions, respectively.

For Elp456 + tubulin samples (Fig. S6A), after gel filtration as described in the “Tubulin binding experiments” section, the complex-containing fractions were pooled and crosslinked using 1.5 mM BS3 (bis(sulfosuccinimidyl)suberate, ThermoFisher, 21580) for 1h on ice. The reaction was quenched by the addition of 50 mM NH_4_HCO_3_. The sample was then vitrified using the same protocol as for Elp456. Data was recorded on a ThermoFisher Scientific Titan Krios electron microscope operating at 300 keV (Fig. S6A) with a calibrated magnification of 96000x, corresponding to a magnified pixel size of 0.824 Å, using a Falcon IV direct electron in counting mode with a dose rate of 5.78 e/Å^2^/s and defocus ranging from -1 to -4 μm. Total exposure time was 8 seconds, resulting in an accumulated dose of ∼ 46 e/Å^2^. EPU software (ThermoFischer Scientific) was used for automated data acquisition. 4420 or 3733 images were collected for the Elp456 and the Elp456+Tubulin samples, respectively.

Movies were imported into RELION 4,0^71,72^. Dose fractionated image stacks were subjected to beam-induced motion correction and filtered according to the exposure dose using RELION’s own implementation. CTFFIND-4.1 was used to estimate the contrast transfer function (CTF) parameters for the motion-corrected micrographs. LoG-based auto-picking was initially used to pick particles from the datasets. Particles were extracted using a box size of 200 Å and, after several rounds of 2D classification (using a 140 Å mask), the best-looking classes were used for template-based picking using RELION’s own implementation. Particles were again extracted and classified using similar parameters as before. Note that several strategies have been tried to favour the visualization of any density that could be attributed to tubulin, including picking techniques, 2D and 3D classification, masking and refinements.

### AlphaFold2 modelling of Drosophila Elp456

The *Drosophila melanogaster* Elp456 model was generated using AlphaFold2 Multimer^40,41^ as implemented in Colabfold^73^. Drosophila melanogaster sequences were provided and identical results were obtained with and without template search in the PDB for the known structures of the Elp456 ring from different speicies^27,28,74^, resulting in both cases in high confidence models. In detail, the core of the six subunits as well as the interfaces were predicted with high confidence and minimal variability between models, whereas the differences were more important in the flexible regions, as expected. The model with the highest overall confidence was retained as the model shown in Sup. Fig. S5E, F and Sup. Fig. S7B, C.

### Cross-linking coupled to mass spectrometry (XL-MS)

For cross-linking coupled mass spectrometry, 75 μl of His-PC-Elongator at 4.2 mg/ml was cross-linked with the N-hydroxysuccinimide (NHS) ester disuccinimidyl dibutyric urea (DSBU). Cross-linking reaction was allowed for 45 min at room temperature with a 150x excess of crosslinker. The reaction was quenched by the addition of NH_4_HCO_3_ to a final concentration of 50 mM, and incubating for 15 min. Note that this experiment was repeated twice using identical conditions (n=2). Cross-linked samples were precipitated with methanol/chloroform^75^ resuspended in 8 M urea, reduced with 10 mM DTT, and alkylated with 50 mM iodoacetamide. Following alkylation, proteins were diluted with 50 mM NH_4_HCO_3_ to a final concentration of 2 M urea and digested with trypsin (Promega), at an enzyme-to-substrate ratio of 1:20, overnight at 37°C or sequentially with trypsin and Glu-C (Promega) at an enzyme-to-substrate ratio of 1:20 and 1:50 at 37°C and 25°C, respectively. The samples were acidified with formic acid to a final concentration of 2% (v/v) then split into two equal amounts for peptide fractionation by peptide size exclusion and reverse phase C18 high pH chromatography (C18-Hi-pH). For peptide size exclusion, a Superdex Peptide 3.2/300 column (GE Healthcare) with 30% (v/v) acetonitrile/0.1% (v/v) TFA as mobile phase and a flow rate of 50 μl/min was used, and fractions collected every two minutes over the elution volume of 1.0−1.7 ml. C18-Hi-pH fractionation was carried out on an Acquity UPLC CSH C18 1.7 μm, 1.0 × 100 mm column (Waters) over a gradient of acetonitrile 2−40% (v/v) and ammonium hydrogen bicarbonate 100 mM.

The fractions were lyophilized and resuspended in 2% (v/v) acetonitrile and 2% (v/v) formic acid and analysed by nanoscale capillary LC-MS/MS using an Ultimate U3000 HPLC (Thermo Fisher Dionex, USA) to deliver a flow of approximately 300 nl/min. A C18 Acclaim PepMap100 5 μm, 100 μm × 20 mm nanoViper (Thermo Fisher Dionex, USA), trapped the peptides before separation on a C18 Acclaim PepMap100 3 μm, 75 μm × 250 mm nanoViper (Thermo Fisher Dionex, USA). Peptides were eluted with a gradient of acetonitrile. The analytical column outlet was directly interfaced via a nano-flow electrospray ionization source, with a hybrid quadrupole orbitrap mass spectrometer (Orbitrap Q-Exactive HF-X, Thermo Scientific). Mass-Spectrometry data were acquired in data-dependent mode. High-resolution full scans (R = 120,000, m/z 350-2,000) were recorded in the Orbitrap followed by higher energy collision dissociation (HCD, stepped collision energy 30 ± 3) of the 10 most intense MS peaks. MS/MS scans (R = 45,000) were acquired with a dynamic exclusion window of 20s being applied.

For data analysis, Xcalibur raw files were converted to MGF format by MSConvert (Proteowizard) and put into MeroX^76,77^. Searches were performed against an ad hoc protein database containing the sequences of the complexes and randomized decoy sequences generated by the software. The following parameters were set for the searches: a maximum number of missed cleavages of three; targeted residues K, S, Y and T; minimum peptide length of five amino acids; variable modifications: carbamidomethyl-Cys (mass shift 57.02146 Da), Met-oxidation (mass shift 15.99491 Da); DSBU modification fragments: 85.05276 Da and 111.03203 Da (precision: 5 ppm MS1 and 10 ppm MS2); false discovery rate cut-off: 5%. Finally, each fragmentation spectrum was manually inspected and validated.

### Statistics & Reproducibility

Unless stated otherwise, measurements are given in mean ± SEM. Statistical analyses were performed using GraphPad Prism 8 with an alpha of 0.05. Normality of variables was verified with Kolmogorov-Smirnov tests. Homoscedasticity of variables was always verified when conducting parametric tests. Post-hoc tests are indicated in their respective figure legends. No statistical method was used to predetermine sample size. The experiments were not randomized. All microtubule dynamics experiments were blindly analysed. No data were excluded from the analyses.

All representative results shown for western-blots, protein purifications, microtubule pelleting assays and microtubule TIRFM experiments (including microtubule binding and microtubule dynamics assay) were performed at least three times independently with equivalent results.

## Acknowledgments

This work was supported by the Medical Research Council, as part of United Kingdom Research and Innovation (UK Research and Innovation; grant no. MC_UP_1201/13 to E.D.) and the Human Frontier Science Program (Career Development Award grant no. CDA00034/2017 to E.D.). VJ.P.-H. was supported by an EMBO postdoctoral fellowship (ALTF 577-2018). K.E.M. is supported by the Wellcome Trust through a Sir Henry Wellcome Postdoctoral Fellowship (220480/Z/20/Z). CJ. was supported by the Agence Nationale de la Recherche ANR-10-1 DEX-0001-02, LabEx Cell(n)Scale ANR-11-LBX-0038, Institut Curie, the French National Research Agency (ANR) awards ANR-17-CE13-0021 and ANR-20-CE13-0011, and the Fondation pour la Recherche Medicale (FRM) grant DEQ20170336756. M.M.M is supported by the Fondation Vaincre Alzheimer FR-16055p and the France Alzheimer grant 2023. M.G. is supported by Institut Curie 3-i PhD Program (IC-3i). We are grateful to the electron microscopy facility staff at the MRC Laboratory of Molecular Biology for access and support for sample preparation and data collection, as well as J. Grimmett, T. Darling and I. Clayson for maintenance of the LMB scientific computing infrastructure. We thank the LMB Biophysics facility for access and maintenance of the MST and C-trap equipment, and specially C. Batters for his help with combined IRM-TIRF imaging. We thank the electronics workshop of the LMB, in particular M. Kyte for his help with the Field Programmable Gate Array hardware required for precise synchronization of our microscope. We also thank C. Lau and S. Chaaban for help with a local implementation of AlphaFold2. We thank S. Bullock for critical reading of the manuscript.

## Author contributions

VJ.P.-H. conducted most experiments in this work, in particular all biochemical experiments, microtubule assays and associated analysis, as well as CryoEM sample preparation, data collection and analysis. A.B. and K.E.M. optimized glass coverslip passivation for TIRFM. K.E.M. optimized the BiGBac vectors. G.D. performed and analysed XL-MS. M.G. and M.M.M. purified mouse and HeLa S3 tubulin. VJ.P.-H., CJ. and E.D. conceptualized the project. All authors contributed to writing the paper.

