## Supplementary Table and Figures for "Elongator is a microtubule polymerase selective for poly-glutamylated tubulin"

### Supplementary Tables

**Supplementary Table 1. Detected crosslinks between Elp123 and Elp456 from a His-PC-SNAP-Elongator prep from *Drosophila* S2 cells.**

| Protein1 | Protein2 | Position 1 | Position 2 | Score |
|---|---|---|---|---|
| dElp1 | dElp1 | 168 | 230 | 53 |
| dElp1 | dElp1 | 896 | 870 | 139 |
| dElp1 | dElp1 | 966 | 915 | 92 |
| dElp1 | dElp1 | 1071 | 696 | 145 |
| dElp1 | dElp1 | 1184 | 930 | 82 |
| dElp1 | dElp1 | 302 | 377 | 136 |
| dElp1 | dElp1 | 696 | 740 | 62 |
| dElp1 | dElp1 | 870 | 896 | 151 |
| dElp1 | dElp1 | 915 | 966 | 126 |
| dElp1 | dElp1 | 930 | 1115 | 79 |
| dElp1 | dElp1 | 930 | 1161 | 124 |
| dElp1 | dElp1 | 930 | 1184 | 136 |
| dElp1 | dElp1 | 1115 | 1235 | 158 |
| dElp1 | His-PC-dElp3S | 854 | 97 | 79 |
| dElp1 | dElp4 | 540 | 256 | 149 |
| dElp1 | dElp4 | 1115 | 364 | 64 |
| dElp1 | dElp6_anon-i1 | 631 | 228 | 187 |
| dElp2 | dElp2 | 97 | 58 | 102 |
| dElp2 | dElp2 | 97 | 756 | 134 |
| dElp2 | dElp2 | 601 | 456 | 127 |
| dElp2 | dElp2 | 756 | 58 | 113 |
| dElp2 | dElp2 | 58 | 97 | 103 |
| His-PC-dElp3S | His-PC-dElp3S | 69 | 106 | 72 |
| His-PC-dElp3S | His-PC-dElp3S | 75 | 61 | 172 |
| His-PC-dElp3S | His-PC-dElp3S | 97 | 60 | 170 |
| His-PC-dElp3S | His-PC-dElp3S | 97 | 106 | 59 |
| His-PC-dElp3S | His-PC-dElp3S | 102 | 60 | 92 |
| His-PC-dElp3S | His-PC-dElp3S | 61 | 75 | 126 |
| His-PC-dElp3S | His-PC-dElp3S | 69 | 106 | 92 |
| His-PC-dElp3S | His-PC-dElp3S | 69 | 419 | 75 |
| His-PC-dElp3S | His-PC-dElp3S | 106 | 360 | 63 |
| dElp4 | dElp4 | 169 | 256 | 75 |
| dElp4 | dElp4 | 276 | 11 | 100 |
| dElp4 | dElp5_poly | 64 | 159 | 130 |

| | | | | |
|---|---|---|---|---|
| dElp4 | dElp5_poly | 364 | 159 | 126 |
| dElp4 | dElp5_poly | 364 | 159 | 99 |
| dElp4 | His-PC-dElp3S | 364 | 360 | 89 |
| dElp4 | His-PC-dElp3S | 364 | 360 | 140 |
| dElp4 | dElp4 | 11 | 276 | 99 |
| dElp5_poly | dElp5_poly | 9 | 159 | 109 |
| dElp5_poly | dElp5_poly | 27 | 175 | 81 |
| dElp5_poly | dElp5_poly | 179 | 151 | 83 |
| dElp5_poly | dElp5_poly | 179 | 151 | 111 |
| dElp5_poly | dElp6_anon-i1 | 27 | 161 | 132 |
| dElp5_poly | His-PC-dElp3S | 151 | 31 | 103 |
| dElp5_poly | His-PC-dElp3S | 151 | 61 | 76 |
| dElp5_poly | His-PC-dElp3S | 151 | 75 | 72 |
| dElp6_anon-i1 | dElp6_anon-i1 | 209 | 228 | 50 |
| dElp6_anon-i1 | dElp6_anon-i1 | 209 | 250 | 113 |
| dElp6_anon-i1 | His-PC-dElp3S | 199 | 419 | 88 |
| dElp6_anon-i1 | His-PC-dElp3S | 209 | 419 | 56 |
| dElp6_anon-i1 | His-PC-dElp3S | 199 | 419 | 187 |
| dElp6_anon-i1 | His-PC-dElp3S | 209 | 419 | 174 |

### Supplementary Figures

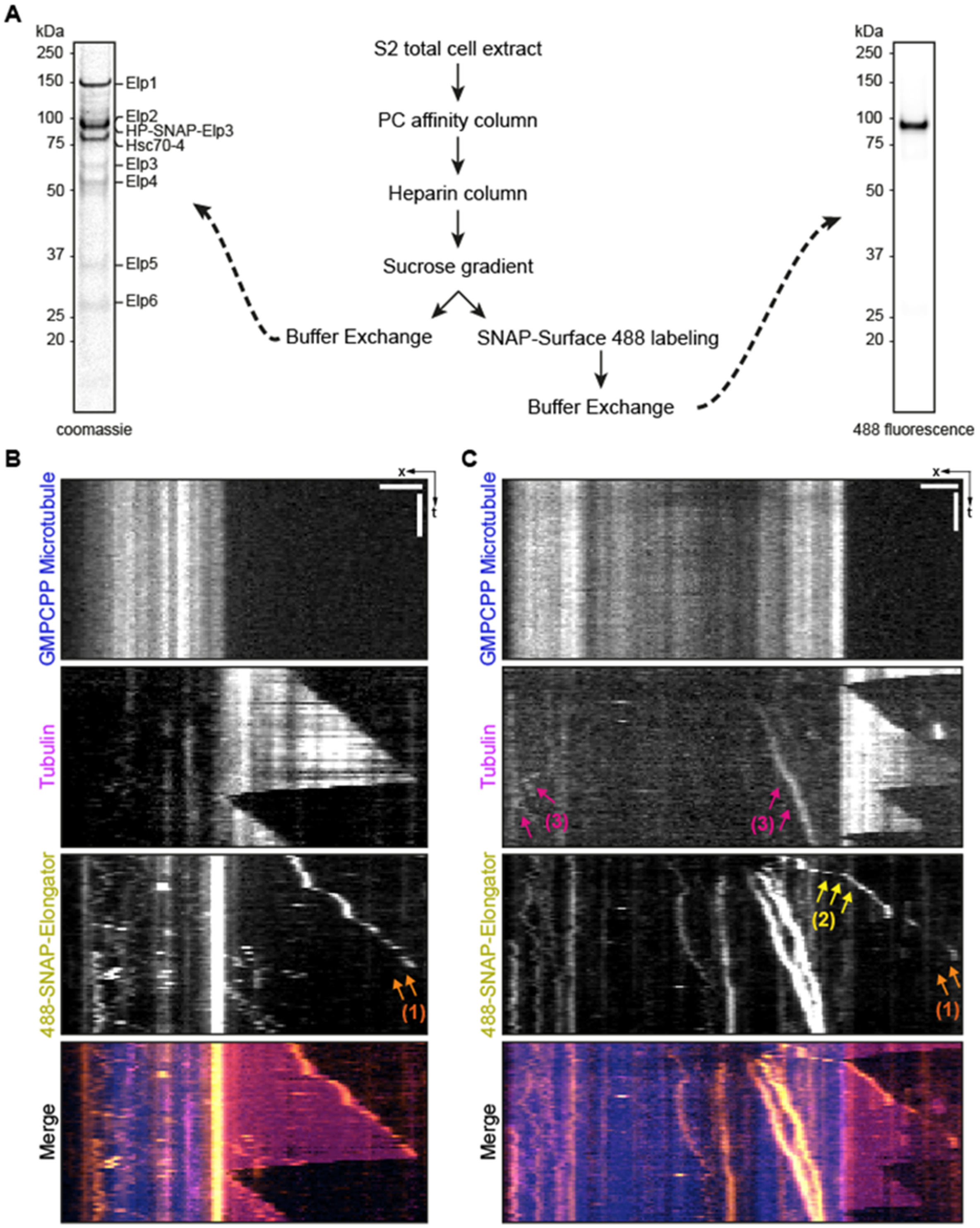


**Supplementary Figure S1. Elongator purification and characterization**

**(A)** Purification and fluorescent labelling of Elongator complex from *Drosophila* S2 cells (see methods). **(B, C)** Biotinylated, rhodamine-labelled GMPCPP-stabilized seeds (red) are anchored via NeutrAvidin to PLL-PEG-Silane. Free tubulin (10% HiLyte 647-labelled, cyan) and Alexa488-SNAP-Elongator is added and Behaviours of Elongator complex observed on microtubules by TIRFM: (1) Elongator detaches from the microtubule ends when microtubules undergo catastrophe (orange arrows); (2) Elongator can "jump" from the GMPCPP seed to the dynamic microtubule, tracking the growing end (yellow arrows); Tubulin signal can be observed diffusing together with Elongator (magenta arrows). Note that 488-Elongator signal can also be observed diffusing on the microtubules.

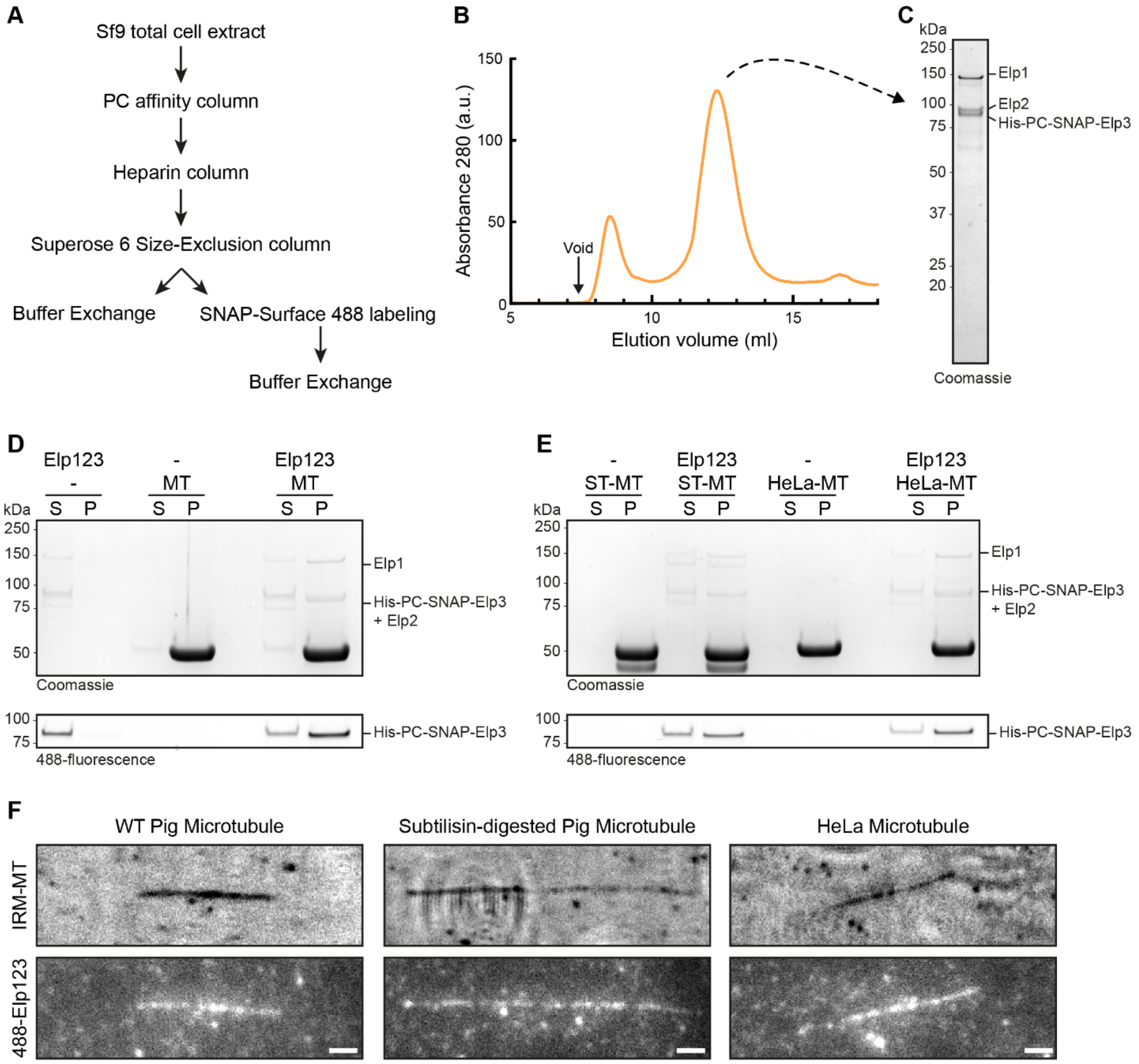


**Supplementary Figure S2. Elp123 purification and characterization**

**(A, B, C)** Purification and fluorescent labelling of Elp123 from Sf9 cells (see methods). **(D, E)** Coomassie (top) and 488-fluorescence (bottom) analysis of *in vitro* co-sedimentation assays with microtubules stabilized with taxol (5 µM tubulin, 40 µM taxol) and 160 nM Elp123. In the presence of pig brain microtubules (D), partially subtilisin-digested pig brain microtubules (E) and HeLa S3 microtubules (E), all Elp123 is found in the pellet. P, pellet; S, supernatant. **(F)** 488-Elp123 decorates wild-type pig microtubules (left), partially subtilisin-digested pig brain microtubules (middle) and HeLa S3 microtubules (right). Scale bars = 2 µm.

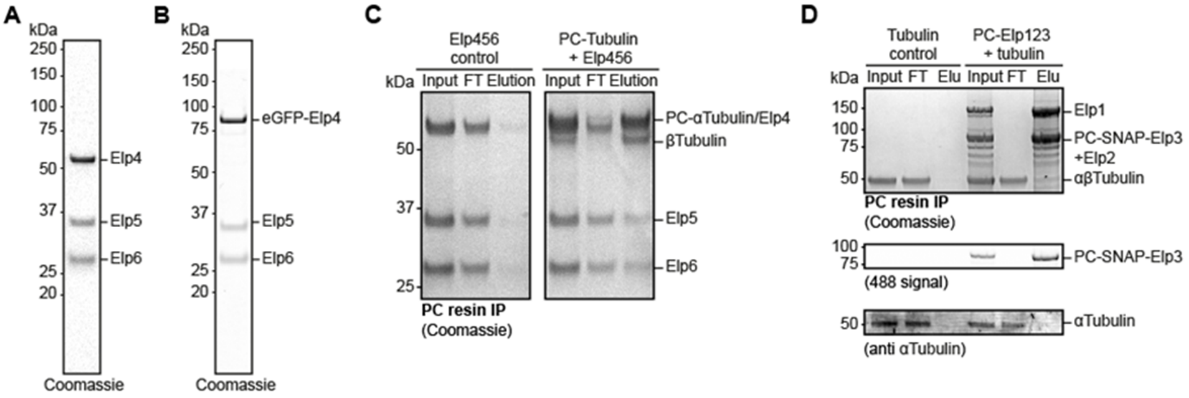


**Supplementary Figure S3. Elongator sub-complexes binding to tubulin**

**(A, B)** Purification of Elp456 and eGFP-Elp456 from *E. coli* cells. **(C)** Elp456 binds to recombinant *Drosophila* PC-tagged α1β1-tubulin heterodimers. Immunoprecipitation assay using a resin coated with anti-PC antibodies. Elp456 is detected in the elution only in the presence of PC-tubulin. Protein presence was analysed using coomassie blue **(D)** Elp123 does not bind to tubulin. Pig brain tubulin cannot be detected eluting from PC-resin in the absence or presence of PC-Elp123. A very high concentration of Elp123 was used to favour any possible weak binding between tubulin and Elp123. The coomassie blue analysis was complemented with fluorescence (488-, labelling Elp123) and western blot analysis using antibodies against αtubulin.

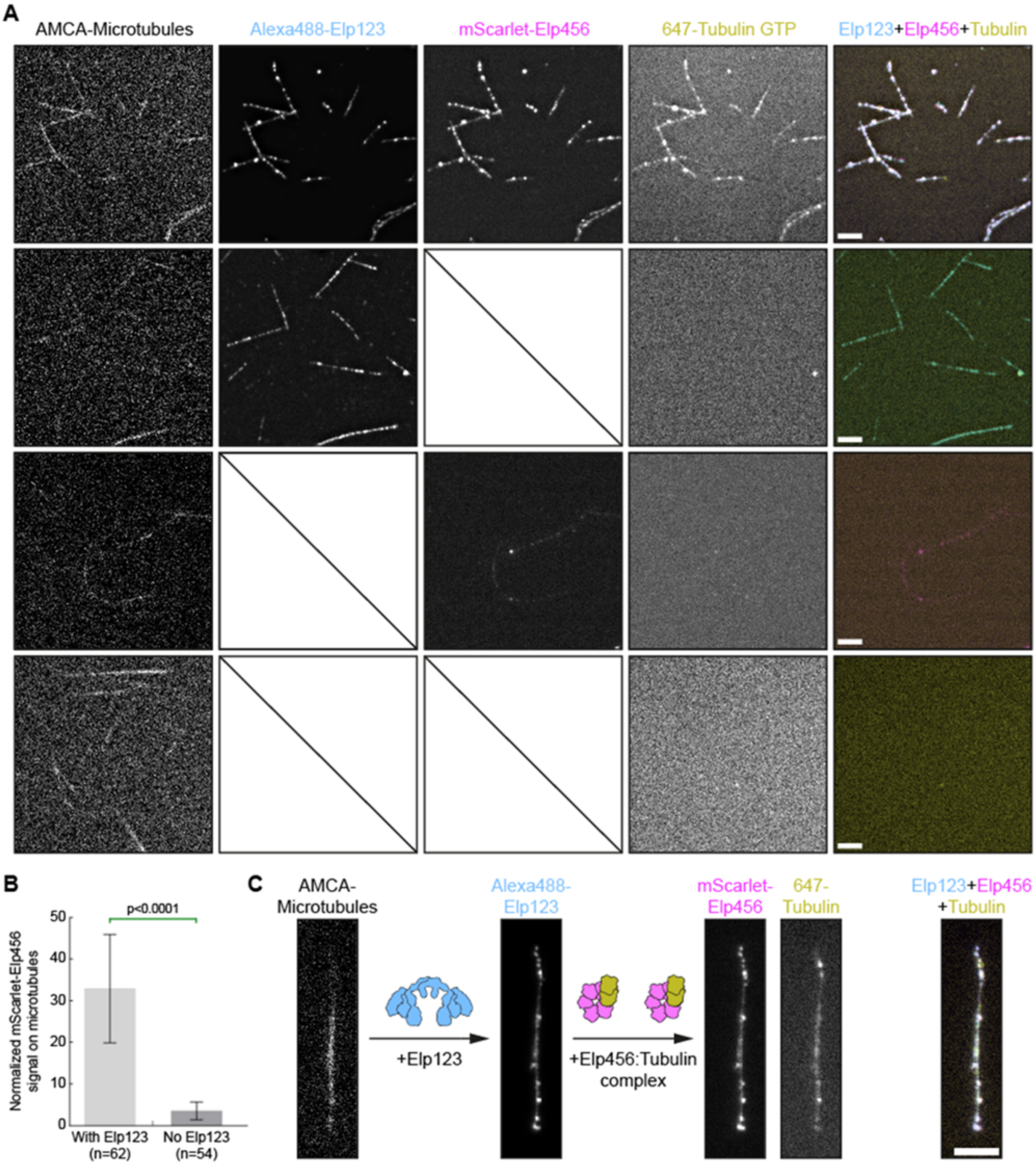


**Supplementary Figure S4. Elongator-tubulin complex reconstitution on microtubules**

**(A)** Controls for Fig. 4. Note that the first row is duplicated from Fig. 4 here for convenience. In the absence of Elp456, no tubulin signal can be detected on microtubules (second row). Similarly, in the absence of both Elp123 and Elp456, no tubulin signal is detected on microtubules (bottom row). Note that in the absence of Elp123, a weak mScarlet-Elp456 signal can be observed on microtubules (third row). This signal is however significantly weaker than when Elp123 is also present ($p<0.0001$ two-tailed, unpaired t-test). **(C)** A pre-formed mScarlet-Elp456:647-Tubulin complex can be recruited to Elp123-decorated microtubules. Scale bars = 5 µm.

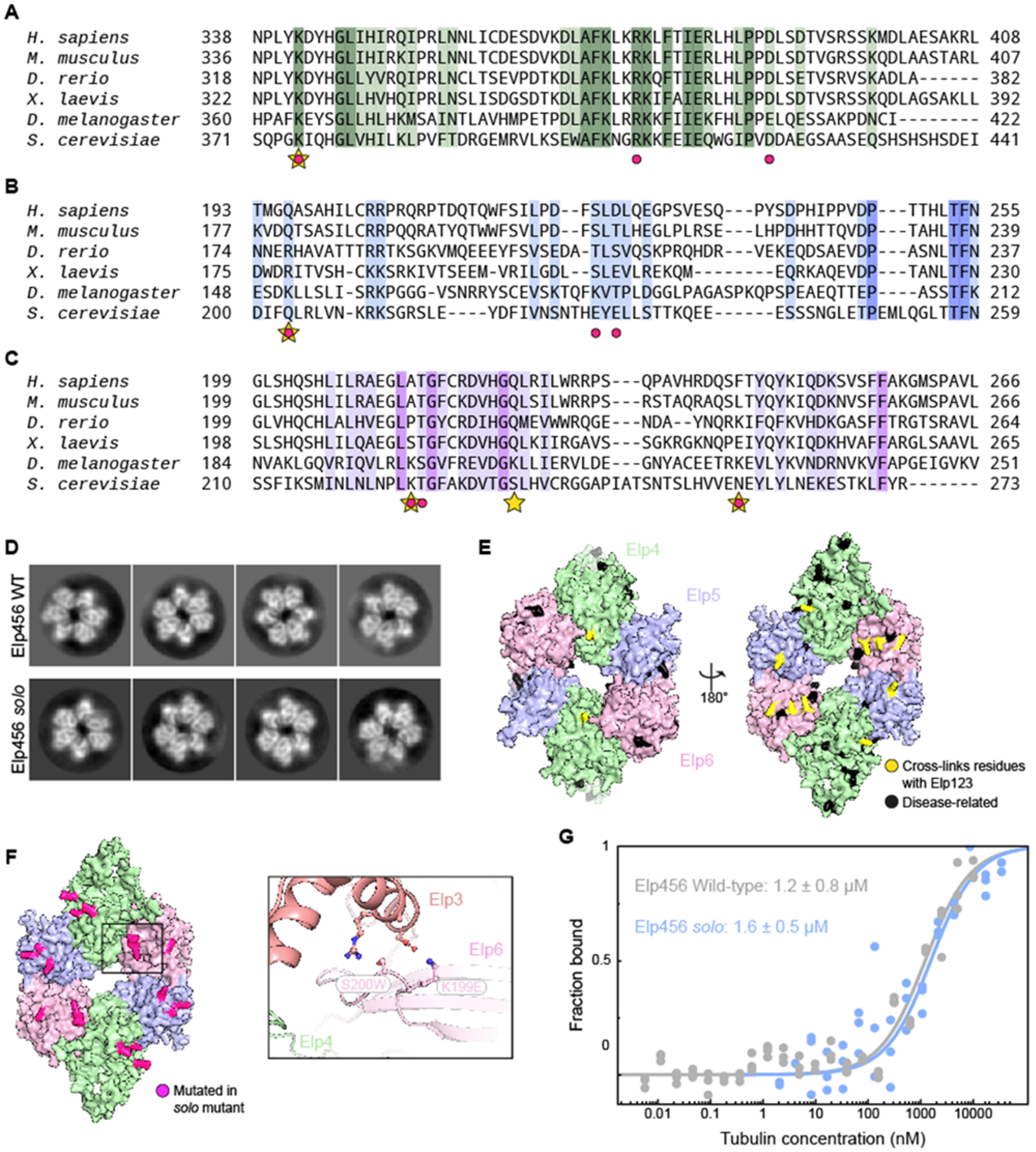


**Supplementary Figure S5. Elongator-tubulin complex reconstitution on microtubules**

**(A, B, C)** Protein sequence alignments of regions of Elp4 (A), Elp5 (B) and Elp6 (C) surrounding detected crosslinks between Elp123 and Elp456 (E) (see also methods). Dark colours: residue strictly conserved. Lighter colours: partially conserved. Residue number and species used for the alignments are indicated. Yellow stars highlight detected crosslinks. Magenta dots indicate residues mutated in Elp456 *solo* mutant: Elp4 K364E, R397E, E410R; Elp5 K151E, K179E, T181A; Elp6 K119E, S200W, K228E. **(D)** 2D class averages of Elp456 WT (top) and Elp456 *solo* mutant (bottom). 140Å mask. **(E, F)** Alphafold model of *Drosophila* *melanogaster* Elp456 (see methods and Sup. Fig. S7B). **(E)** Detected crosslinks with Elp123 are highlighted in yellow. Disease-related residues found in ActiveDriverDB database. **(F)** Mutated residues in Elp456 *solo* mutant. Inset shows the interface between Elp3 and Elp6 from ref[1], highlighting how the mutations K199E and S200W introduced in Elp456 *solo* might perturb this interaction. **(G)** Elp456 *solo binds* to recombinant *Drosophila* α1β1-tubulin heterodimers with wild-type affinity. Calculated dissociation constant ($K_d$) values are indicated (mean ± s.d.; n=3).

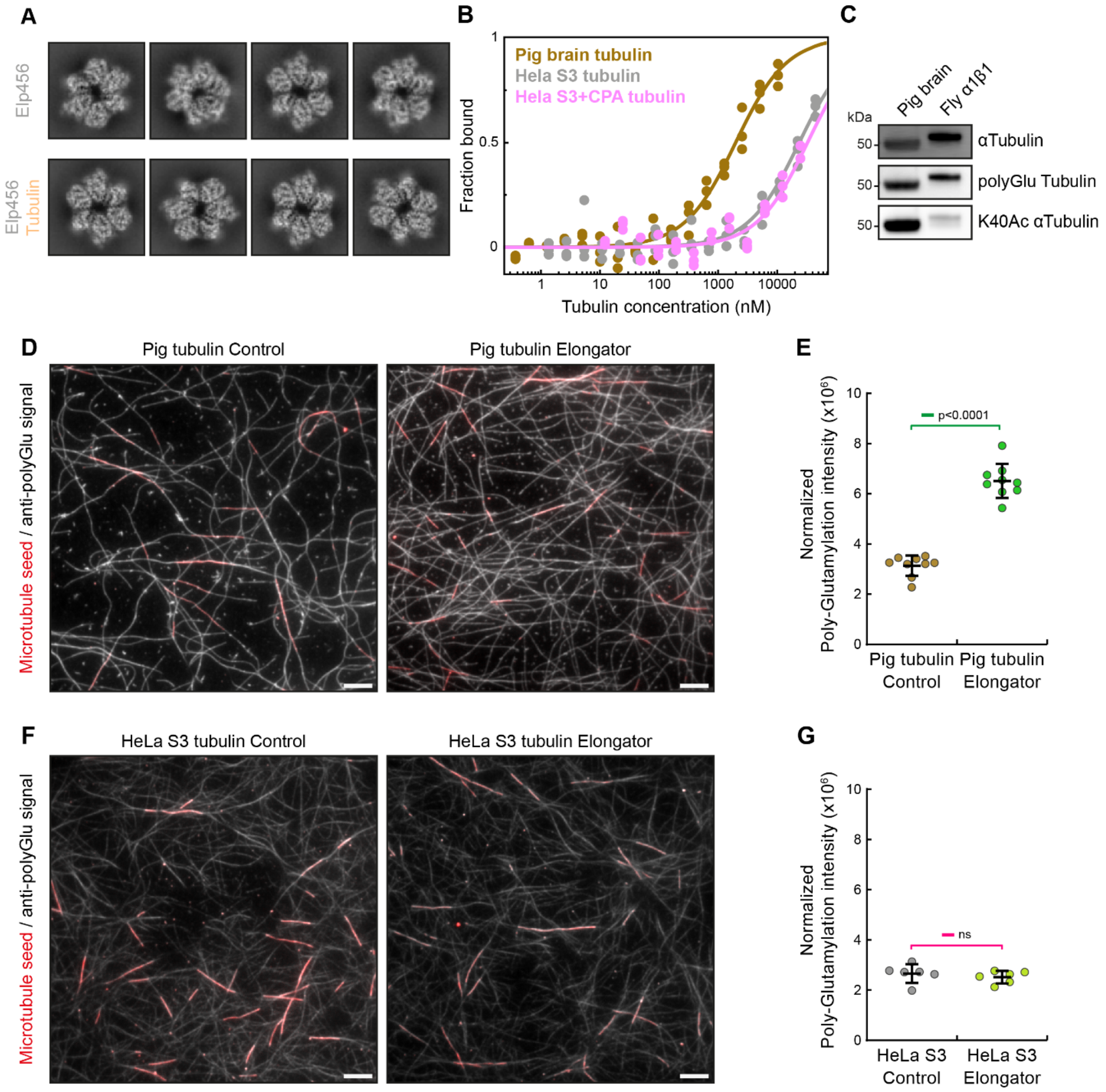


**Supplementary Figure S6. Elongator is a poly-glutamylated tubulin polymerase.**

**(A)** 2D class averages of Elp456 WT (top) and Elp456:tubulin complex isolated from size exclusion chromatography (as in Fig. 3B) and crosslinked with 1.6 mM BS3 for 1h. No tubulin extra density can be detected in the presence of tubulin (see methods). **(B)** Measurement of the eGFP-Elp456 and αβ-tubulin heterodimers (as indicated) interaction using microscale thermophoresis. Calculated dissociation constant ($K_d$) values are indicated (mean ± s.d.; n=3). See also Fig. 6A, B. **(C, E)** Representative field of view of microtubules labelled with fluorescently-labelled anti-polyglutamylated tubulin antibodies (gray) and HiLyte488-labelled GMPCPP stable seeds (red). Conditions as indicated. **(D, F)** Quantification of total poly-glutamylated tubulin signal (see methods). Scale bars = 5 µm.

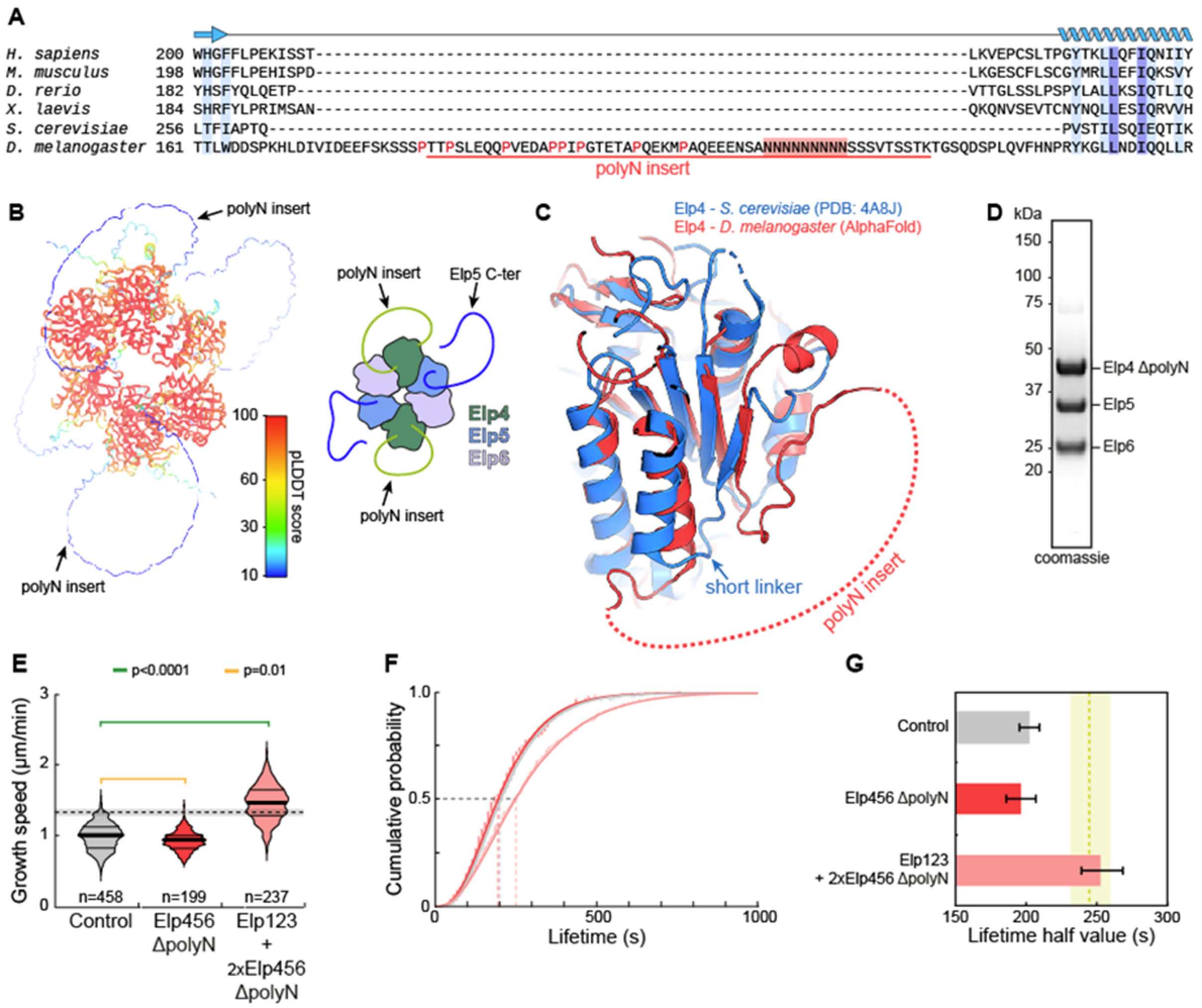


**Supplementary Figure S7. A specific *D. melanogaster* insert in Elp4 does not participate in microtubule stabilization**

**(A)** Protein sequence alignment of Elp4 surrounding a specific insert present only in *D. melanogaster* (polyN insert). Residue number and species used for the alignments are indicated. Indicated are known structural elements of Elp4 (top). Dark blue: residue strictly conserved. Light blue: partially conserved. Residue number and species used for the alignments are indicated. Also indicated is the enrichment in proline (red contour) and asparagine (red background) residues in the insert. The red line indicates the residues removed in the Elp456 ΔpolyN mutant. **(B)** Alphafold model of *D. melanogaster* Elp456. pLDDT score is indicated. Note how the polyN insert is predicted to be disordered and facing the solvent, making it a perfect candidate for protein-protein interactions. Elp5's C-terminal tail is similarly predicted. A simplified drawing is shown on the right. **(C)** Elp4 superposition between the yeast (PDB 4A8J) and the fly (AlphaFold) subunits highlighting the location of the polyN insert. **(D)** Purification of Elp456 ΔpolyN mutant. **(E, F, G)** Effect of the indicated conditions on the growth speed (E) and lifetime (F, G) of microtubules at the plus end. n, number of microtubule-growing events analysed. **(E)** P values for a Kruskal-Wallis test followed by Dunn's multiple comparison test are indicated. Dashed line represents an increase of ~1.4 in the speed of microtubule growth[2]. Thick line, median; thin line, quartile. **(F, G)** Lifetime estimate ± error from the bootstrapped mean lifetimes (see methods) are indicated in the right panel.

Green shaded region indicates the values obtained with wild-type Elp456 (Fig. 4C). Dashed line represents the median, and the shaded region the quartiles.
